# Breaking a Strong Amide Bond: Structure and Properties of Dimethylformamidase

**DOI:** 10.1101/2019.12.17.879908

**Authors:** Chetan Kumar Arya, Swati Yadav, Jonathan Fine, Ana Casanal, Gaurav Chopra, Gurunath Ramanathan, Kutti R. Vinothkumar, Ramaswamy Subramanian

## Abstract

Dimethylformamidase (DMFase) breaks down the human-made synthetic solvent *N,N*-dimethyl formamide(DMF) used extensively in industry(1). DMF is not known to exist in nature and was first synthesized in 1893. In spite of the recent origin of DMF, certain bacterial species such as *Paracoccus, Pseudomonas*, and *Alcaligenes* have evolved pathways to breakdown DMF and use them as carbon and nitrogen source for growth(2, 3). The work presented here provides a molecular basis for the ability of DMFase from *Paracoccus* to function in exacting conditions of high solvent concentrations, temperature and ionic strength to catalyze the hydrolysis of a stable amide bond. The structure reveals a multimeric complex of the α_2_β_2_ type or (α_2_β_2_)_2_ type. One of the three domains of the large subunit and the small subunit are hitherto undescribed folds and as yet of unknown evolutionary origin. The active site is made of a distinctive mononuclear iron that is coordinated by two tyrosine residues and a glutamic acid residue. The hydrolytic cleavage of the amide bond is catalyzed at the Fe^3+^ site with a proximal glutamate probably acting as the base. The change in the quaternary structure is salt dependent with high salt resulting in the larger oligomeric state. Kinetic characterization reveals an enzyme that shows cooperativity between subunits and the structure provides clues on the interconnection between the active sites.

**Significance Statement:** *N,N*-dimethyl formamide(DMF) is a commonly used industrial solvent that was first synthesized in 1893. The properties that make DMF a highly desired solvent also makes it a difficult compound to breakdown. Yet, certain bacteria have evolved to survive in environments polluted by DMF and have enzymes that breakdown DMF and use it as their carbon and nitrogen source. The molecular structure of the enzyme that breaks down the stable amide bond in these bacteria, reveals two new protein folds and a unique mononuclear iron active site. The work reported here provides the structural and biochemical framework to query the evolutionary origins of the protein, as well as in engineering this enzyme for use in bioremediation of a human made toxic solvent.

## INTRODUCTION

Dimethylformamide (DMF) is an organic solvent commonly used in chemical synthesis, leather, printing, and petrochemical industries. The extent of its usage is evident from the quantity of its production(4, 5). Its polar nature accords properties similar to water, and its physicochemical features make it versatile(miscible in many organic solvents and water, a high boiling point of 153 °C, etc.) (1, 6–8). It is a known hepatotoxic and ecotoxic agent(9, 10). DMF is remarkably resilient to chemical and photochemical decomposition. Biodegradation often emerges as an efficient process for its removal from the environment(11). In spite of the short history of DMF on earth (introduced in 1893), several microbes that can grow using DMF as the sole carbon source have been described, with most of them belonging to the phylum Proteobacteria (*Pseudomonas, Alcaligenes, Orchobactrum* sp., and *Paracoccus* species) (2, 3, 12–14). *Pseudomonas* uses an oxidative demethylation pathway to convert DMF into formamide. The product is subsequently converted into ammonia and formate by the enzyme formamidase. Other organisms encode a *N, N*-dimethylformamidase (DMFase) that catalyzes the formation of formate and dimethylamine(15). DMFases from different organisms have been characterized, and all of them comprise two polypeptide chains, a smaller polypeptide of 15 kDa, and a larger polypeptide of 85 kDa. Sequence-based homology searches do not show any similarity to other ubiquitous amidohydrolases, and the nature of its active site remained a puzzle. Here, we report structures of DMFase determined by electron cryo-microscopy and X-ray crystallography augmented with its biochemical characterization. Enzyme kinetic measurements with different substrates complement the structural data. Based on the structure, we performed site-directed mutagenesis of specific residues to test their role in metal binding and catalysis. Together, these data allow us to propose a plausible mechanism and a molecular explanation for the narrow substrate specificity of the enzyme.

## RESULTS

### Electron cryo-microscopy

When purified native or recombinant DMFase was imaged on ice by electron cryo-microscopy (cryoEM), the micrographs unambiguously revealed two populations in solution (Supplementary Data Fig. 1A). A smaller unit measuring ∼100Å diameter and a larger structure ∼150Å in diameter could be identified. Subsequent 2D classification showed that the smaller unit was a dimer and the larger structure was a dimer of dimers. The dimer of DMFase is made of two α and two β subunits with a total mass of 200 kDa. The dimer of dimers (called tetramer here) has a total mass of 400 kDa.

**Figure 1:**
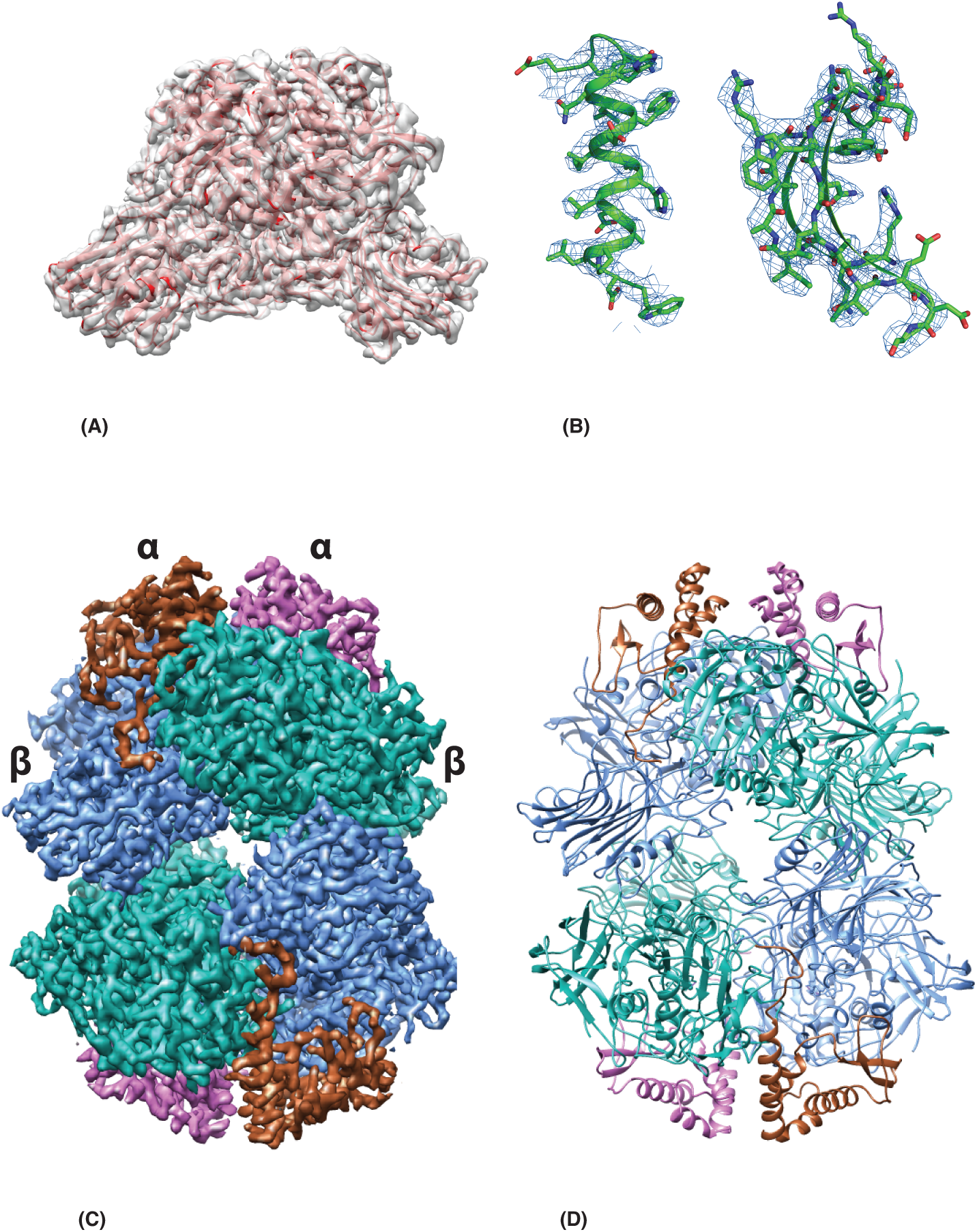
(A) The initial averaged cryo-EM map of the α_2_β_2_ structure with a trace of the main chain. (B) Representative regions of the electron density map at ∼3Å resolution that allowed tracing of the entire chain. (C) Tertiary structure of the tetrameric form made of two dimer of dimers. The α and β subunit of one of the α_2_β_2_ is shown. In brown and purple are the smaller subunits and in blue and green the large subunits. (D) A cartoon diagram of the tetrameric model.

The initial data we collected had a mixture of populations (Supplementary Data Fig.1 A & B), and the reconstructions resulted in a 3.4 Å map for the dimer and a 3.8 Å map for the tetramer. The two populations observed by cryoEM were intriguing. We noted that the salt concentration in the buffer shifts the oligomer (dimer-tetramer) equilibrium. In the presence of low salt, the enzyme is predominantly in the dimeric form (Supplementary Data Fig. 1 C & D), and this equilibrium shifted to the tetramer at higher salt concentrations (≥200 mM) (Supplementary Data Fig.1 E & F). Subsequently, we collected two data sets with no salt and 200 mM NaCl yielding reconstructions with an overall resolution of 3Å (dimer) and 2.8 Å (tetramer), respectively (Fig. 1A, 1C, Supplementary Data Fig. 2, 3). The local resolution plot of both maps showed that much of the reconstruction was resolved between 2.5-3.5 Å (Supplementary Data Fig. 2&3). The high quality of the maps allowed *de novo* tracing of polypeptide chains of both the large and small subunits (Fig. 1B). We used the dimer map for the initial tracing, and the fit of the model to map is shown in Fig. 1A and 1B. The model from the dimer was used to obtain the tetrameric model (Fig. 1C&D and Supplementary Data Fig. 3, Table 1). The interface between the two dimers in the tetramer is solely through the large subunit (Fig. 1C and 1D).

**Table 1.** Cryo-EM Data collection and refinement statistics for DMFase.

| | DMFase<br>200 kDa,<br>$\alpha_2\beta_2$ | DMFase<br>400 kDa,<br>$2\times(\alpha_2\beta_2)$ | DMFase<br>400 kDa,<br>$2\times(\alpha_2\beta_2)$,<br>Y440A mutant | DMFase<br>400 kDa,<br>$2\times(\alpha_2\beta_2)$,<br>E521A mutant |
| --- | --- | --- | --- | --- |
| Data Collection |  |  |  |  |
| Pixel Size (Å) | 1.07 | 1.07 | 1.07 | 1.07 |
| Dose rate (e/p/s) | 0.5 | 0.5 | 0.5 | 0.5 |
| Total dose (e/Å<sup>2</sup>) | 27 | 27 | 27.5 | 27.5 |
| No. of particles | 80674 | 59191 | 18645 | 45851 |
| Resolution (nominal), Å | 3.0 | 2.8 | 3.2 | 3.1 |
| B-factor (sharpening) | 95 | 86 | 95 | 92 |
| Model composition |  |  |  |  |
| Number of chains | 4 | 8 | 8 | 8 |
| Protein (No. of residues) | 1772 | 3544 | 3468 | 3480 |
| Ligand | 2 | 4 | 0 | 0 |
| Model Refinement |  |  |  |  |
| Resolution (Å) | 3.0 | 2.8 | 3.2 | 3.1 |
| Average B-factor |  |  |  |  |
| Overall | 41.7 | 38.8 | 54.7 | 50.6 |
| Protein | 41.7 | 38.8 | 54.7 | 50.6 |
| Ligand (Fe) | 44.2 | 43.1 | - | - |
| RMS deviations |  |  |  |  |
| Bonds (Å) | 0.009 | 0.009 | 0.009 | 0.01 |
| Angles (Å) | 1.2 | 1.2 | 1.3 | 1.3 |
| Validation |  |  |  |  |
| Ramachandram plot<br>(favoured/outliers) | 94.4/0.0 | 95.0/0 | 93.7/0 | 93.6/0 |
| clashscore | 3.4 | 2.9 | 3.4 | 3.3 |
| Molprobity score | 1.5 | 1.4 | 1.5 | 1.5 |

**Figure 2:**
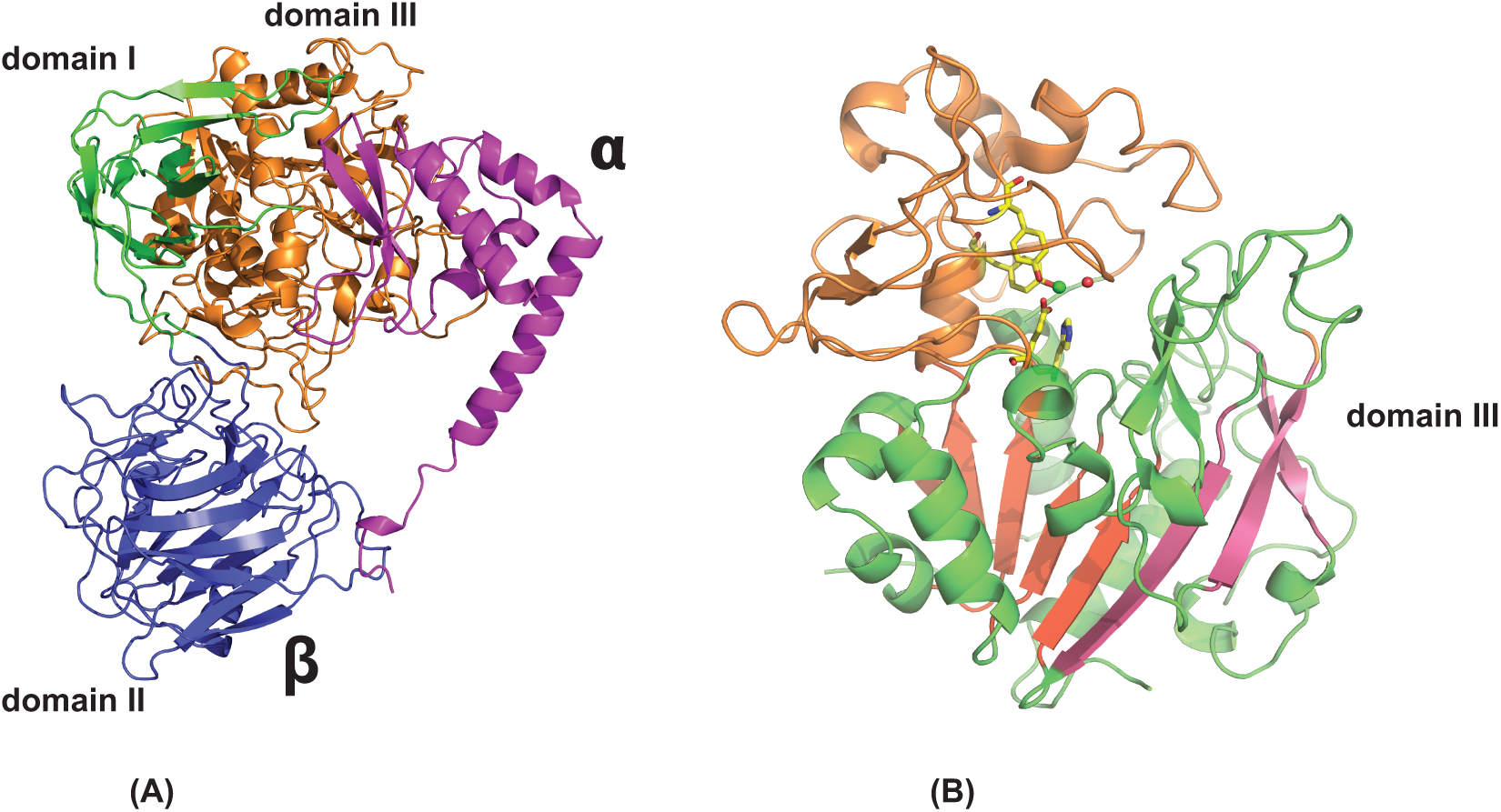
(A) The structure of the unique polypeptide units that forms the smallest structural unit, the αβ hetero-dimer. The small subunit is in purple color and the three domains of the large subunit are in green, orange-brown and blue color. (B) Domain III is shown with the active site. The five parallel beta strands are in orange. The active site residues are in stick representation and the Fe and the liganded water molecule are shown as spheres.

**Figure 3:**
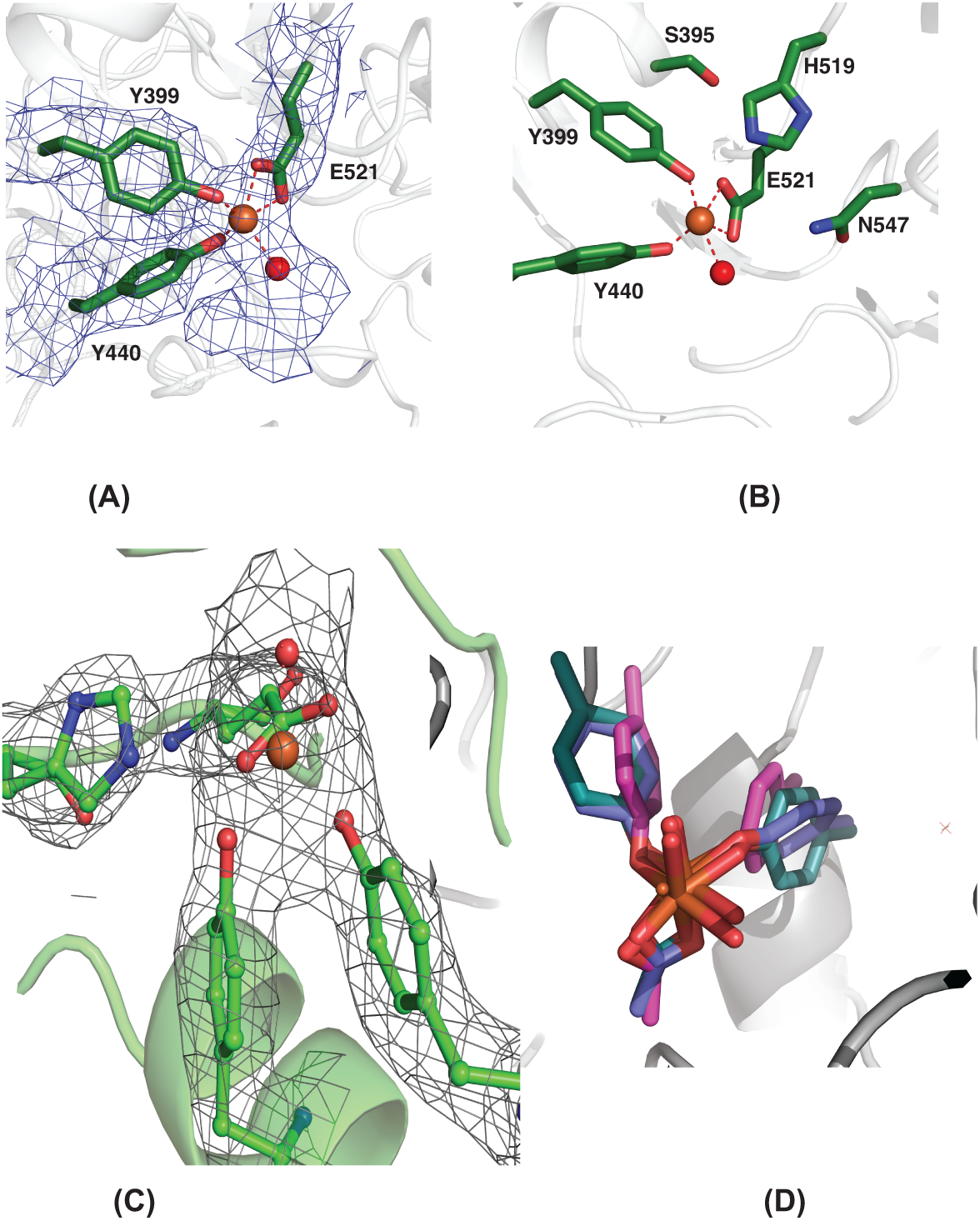
(A) The cryo-EM map around the active site with the modeled Fe, ligands and a single water molecule. (B) The residues in the active site including the essential His 519. (C) The active site residues modeled into the non-crystallographic symmetry-averaged simulated annealed omit map from the crystallographic data. (D) The Fe site modeled from the purple orientations (found from the structures determined) and the relaxed orientations of the ligands after minimization by DFT. The significant rearrangement of the Tyrosine suggests the strain in the active site.

### Crystallography

Subsequently, we determined a crystal structure of DMFase by molecular replacement using the coordinates from cryoEM data. The crystals belong to the space group P2_1_ with two tetramers in the asymmetric unit and the model refined to a resolution of 2.8 Å (Table 2). The structures determined by cryoEM and X-ray crystallography are similar, with the differences observed being minor largely localized to the loop regions.

**Table 2.** X-ray Data Collection and refinement statistics of DMFase.

|  |  |
| --- | --- |
| Wavelength | 1.005 |
| Resolution range | 49.4 - 2.8 (2.85 - 2.8) |
| Space group | P 1 21 1 |
| Unit cell | 160.51 142.55 181.49<br>90 92.9 90 |
| Total reflections | 940355 (48255) |
| Unique reflections | 200660 (9942) |
| Multiplicity | 4.7 (4.9) |
| Completeness (%) | 99.9 (100) |
| Mean I/sigma(I) | 4.4 (1.1) |
| Wilson B-factor | 21.4 |
| R-merge | 0.36 (1.3) |
| R-pim | 0.21 (0.79) |
| CC1/2 | 0.92 (0.32) |
| R-work (R-free) | 0.25(0.28) |
| Number of non-hydrogen atoms | 56520 |
| RMS(bonds)Protein residues | 0.003 |
| RMS(angles) Protein residues | 0.56 |
| Ramachandran favored (%) | 93.38 |
| Ramachandran allowed (%) | 6.28 |
| Ramachandran outliers (%) | 0.34 |
| Rotamer outliers (%) | 5.4 |
| Clash score | 6.85 |
| Average B-factor | 26.45 |
| Number of TLS groups | 16 |

### Two New Folds

All descriptions of the structure described here are based on the best cryo-EM model. The smallest unit is an αβ dimer (Fig. 2A). The large subunit consists of three different domains (Fig. 2B). Using PDBeFOLD(16), we found that domain I is made of residues 1-73 and 338-383 and it adopts an immunoglobulin (IgG)-like fold. Domain II is made of residues 74-337 and adopts the Pentraxin fold. Domain III comprises residues 384-761 and a fold that is not identified by the PDBeFOLD or DALI servers (Fig. 2B). The core of the Domain III fold itself can be described as α/β/α fold, with five β-strands that are parallel and sandwiched between α-helices. Such an arrangement is seen in ThuA like family of proteins. These arrangements are classified as part of the larger family of glutamine amidotransferases(17). The five conserved β-strands, plus the three extra β-strands together form a sheet that forms the inside of the sandwich (Fig. 2B). The connecting region between the β-strands forms a sub-domain that is made of four anti-parallel β-strands that cap the structure. Domains I and III show a significant number of interactions, but the interaction of domain II with other domains of the monomer is somewhat limited (Fig. 2A).

The small subunit of DMFase consists of four α-helices and two β-strands. There are two long helices at the N and C-terminus, with the N-terminal helix wedging into the interface between the two large subunits (Fig. 1C & 1D). As a whole, the small subunit is also an undescribed fold. However, the two helices alone superpose very well with the two helices of ESCRT-1(18) (PDB-ID 2F66). Most of the interactions of the small subunit are with domain III of the large subunit (Fig. 2A). The *N*-terminal residues of the small subunit interact with domain II and seem to stabilize the tertiary structure by holding this domain from adopting other orientations with respect to domains I and III (Fig. 2A). The large subunit, when expressed and purified without the small unit, was not enzymatically active (Supplementary Data Table 1). The size exclusion chromatography profile suggests that it most likely exists as a dimer, albeit with a lower T_m_ (46°C) than the α_2_β_2_ enzyme (Supplementary Data Fig. 4). Thus, the small subunit is most likely to play a role in structural stabilization.

**Figure 4:**
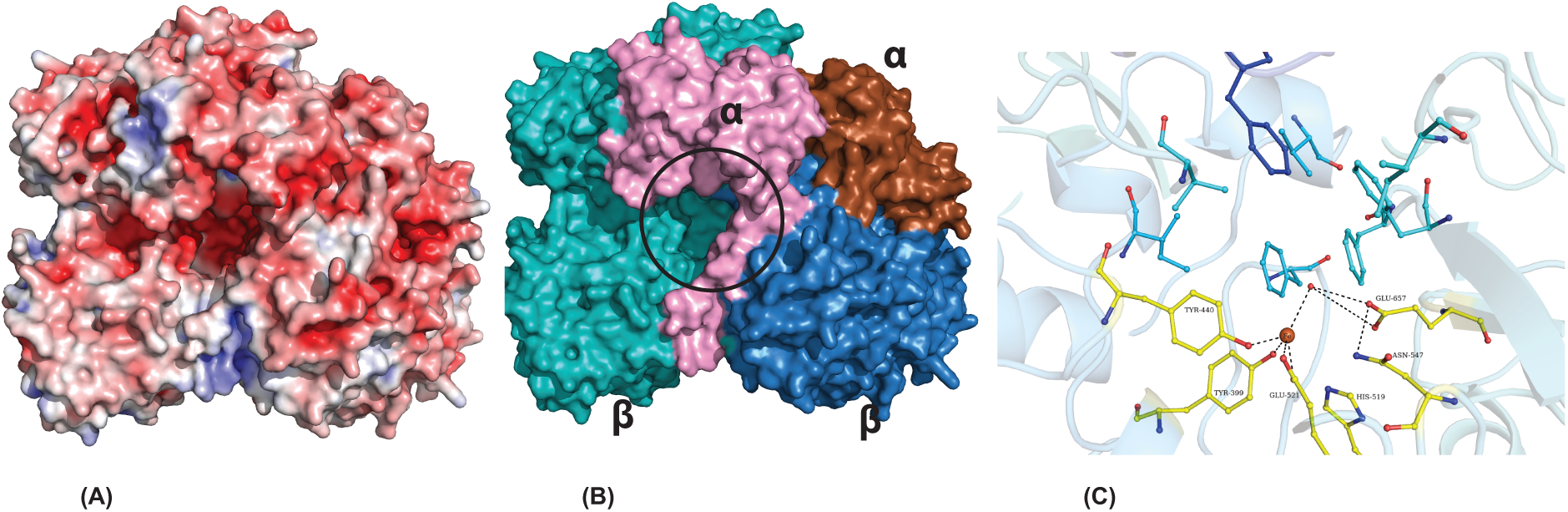
(A) Surface charge distribution calculated by APBS showing the entrance to the active site. (B) Same view as (A), but the subunits of the dimer are colored differently. The entrance to the active site (shown in black circle) has residues from both the large and small subunits. (C) A closer look at the binding pocked. The ligands to the Fe^+3^ and the residues involved in catalysis are labeled. The substrate binding site is made of a number of bulky aromatic residues blueish green. In dark blue is Phe 693 from the loop of the neighboring large subunit.

### A Distinctive Metal Site

The initial EM map clearly showed a large density around residues Y440, Y399, and E521 that could not be accounted for by amino acids (Fig. 3A). We predicted this to be the active site, which is buried in domain III. The exact nature of the metal ion as Fe was confirmed by X-ray fluorescence spectroscopy at the synchrotron (Supplementary Data Fig. 5). EPR spectra collected on the *wild-type* protein suggested that under the conditions of purification and storage of the protein, the iron is mostly in the Fe^3+^ high-spin state. The mononuclear Fe atom is tetracoordinated. It is held by two phenolic hydroxyls of the tyrosine residues (Y399 and Y440 serve as monodentate ligands), while a glutamic carboxylate side chain (E521 serves as a bidentate ligand) (Fig. 3B). A histidine residue (H519) is observed close to the active site (Fig. 3B, 3C). However, in the model derived from both EM and X-ray maps, this histidine residue does not coordinate with the Fe^3+^ ion in the active site. The distance between the Nε2 atom of H519 and Fe^3+^ is approximately 4.0Å. When residues Y440 and E521 were mutated to alanine, the resultant mutants were found to be catalytically inactive with a significantly reduced thermal stability (Supplementary Data Fig. 6; Supplementary Data Table 1). Structures of these mutant enzymes determined by cryoEM show the loss of Fe atoms (Supplementary Data Fig. 7; Table 1). The absence of Fe in these proteins results in the interface region between the large and small subunits becoming disordered (Supplementary Data Fig. 8). We also mutated residues H519 and S395, which are close to the metal-binding center. Of these, S395A showed comparable activity to wild-type DMFase, but the H519A mutant was catalytically inactive (Supplementary Data Table 1). Despite not coordinating Fe^3+^, H519 still plays a role in catalysis.

The coordination of Fe^3+^ is strained and is close to a square pyramidal geometry with two vacant sites (in the absence of modeled water). There is density at the active site that can easily accommodate at least one water molecule in both the EM maps and the crystallographic map, which has been currently modeled (Fig. 3A&C). There are also some differences in the orientation of the residues in the active site in the final refined models of the EM and X-ray. However, the limited resolution does not allow us to interpret these differences with sufficient confidence. In the X-ray and EM maps, there remains unexplained additional density after the addition of Fe and water (Fig. 3A), where common buffer components such as phosphate, oxalate, and DMF can be modeled. In the absence of other experimental evidence, we chose not to over-interpret and left it unmodeled.

There are utmost 46 entries in the PDB, where tyrosines act as coordinating ligands to a mononuclear Fe atom. The architecture of the two tyrosines binding to the Fe^3+^ ion is similar to that present in the structure of the ferric binding proteins from *Nisseria gonorrhoeae* (PDB ID 1d9y) and *Yersinia enterocolitica* (PDB ID 1xvx)(19). Both of these also have a histidine coordinating the ligand and are not catalytically active. The architecture with two tyrosine hydroxyls and a glutamic acid side chain carboxylate as observed here, is unique.

In order to confirm that the chemical environment around the iron atom is strained, we calculated the electronic energy of the iron atom and its surrounding environment using the experimentally observed structure (Fig. 3A) as the starting model. We compared this strain energy to the energy of a relaxed system (Fig. 3D). Electronic energies were calculated using density functional theory (DFT) with the B3LYP functional^20–22^ and the 6-311G(d) basis set in Gaussian16^23^ (Supplementary Data Fig. 9). The difference in the energies obtained between the strained and relaxed octahedral states with Fe^3+^ ions is −222.01 kJ/mol (Supplementary Data Table 2). The difference in energy is significant as the heat of formation for liquid *N,N*-dimethylformamide (DMF) is 239.4±1.2 kJ/mol^24^ and the difference in energy between DMF and the first transition state of decomposition is 27 kJ/mol, suggesting that the relief of this strain is adequate for catalyzing the decomposition of DMF. The specific properties of this unique Fe^3+^ site need further exploration, especially in the context of the reaction mechanism, where a very stable amide bond is hydrolytically cleaved.

### Enzyme Kinetics and Stability

We further characterized this enzyme using a number of biophysical and biochemical tools. Using isothermal calorimetry (ITC), the real-time enzyme kinetics of both native and recombinant DMFase for the substrate DMF were similar (Supplementary Data Fig. 10, 11 and Supplementary Data Table 3). The data from the experiments are a better fit to the Hill equation (with a Hill coefficient of 2) rather than to a classical Michaelis-Menten equation, suggesting cooperativity between subunits (Supplementary Data Fig. 12). Using steady-state kinetics, we show that enzymes purified from *Paracoccus* as well as the recombinantly expressed proteins show similar V_max_ and K_m_ to the DMFase from other organisms(14),(12), (20). A thermal shift assay revealed an optimum T_m_ of DMFase to be 64°C (Supplementary Data Fig. 13) in 250 mM NaCl, revealing that it is a thermotolerant enzyme. The peak activity of *Paracoccus* sourced enzyme was 54°C (Supplementary Data Fig. 14). The temperature dependence of the kinetic and thermodynamic parameters shows reduced enzymatic activity at low temperatures (≤20°C). We then tested the halo-tolerant nature of DMFase by measuring the enzyme activity(21) at increasing salt concentrations. Optimal hydrolase activity for DMFase was detected at 2.5M NaCl. At this salt concentration, the temperature of inversion (T_*i*_) is 56°C (Supplementary Data Table 4). APBS(22) calculation suggests a highly charged surface (Fig. 4A and Supplementary Data Fig. 15), which has been implicated in increased halo-stability in other proteins(23, 24). The α_2_β_2_ buries a total surface area of approximately 9540 Å^2^. The tetramer will have double this surface area buried plus the 1830 Å^2^ at the interface. Thus, one can infer that the large polypeptide and its ability to form oligomers in increased salt concentrations constitute an adaptation that stymies protein denaturation contributing to halostability. When the activity and stability of DMFase was checked at different concentrations of DMF, the enzyme was most active at 0.2M DMF (Supplementary Data Fig. 16 and Supplementary Data Table 5). The enzyme seems to have evolved to work best at 54°C, 1.5M salt, and 0.2M DMF *in vitro*, possibly similar conditions in which the bacterium survives in the sludge.

The substrate specificity of DMFase was tested against compounds such as formamide, acetamide, *N*-methylformamide (NMF), *N*-ethylformamide (NEF), dimethylacetamide (DMAc), benzamide, hexanamide, and urea were studied using real-time ^1^H NMR based kinetic assays. As expected, the enzyme exhibits the highest Vmax for DMF. Formamide, NEF, and NMF were also hydrolyzed with the highest rate observed for NMF but other substrates yielded no detectable products (Supplementary Data Fig. 17, Supplementary Data Table 6).

## DISCUSSION

The structure and experiments reported here provide insights into residues involved in substrate binding and catalysis. A large cavity is observed at the interface between the large and small subunits, which leads towards the active site (Fig. 4B). In the EM-structures of the active site mutants (Y440A and E251A), the interface between the large and the small subunit becomes disordered, indirectly eluding a role for small subunit in the substrate entry pathway (Supplementary Data Fig. 8). The substrate-binding pocket in the active site is constricted by bulky side chains consisting mostly of aromatic residues (Fig 4C). The cavity formed by the hydrophobic residues is large enough to accommodate substitutions other than the dimethyl group on the amine. Currently, there is no structure of the DMF bound form of the protein. Given the nature of the reaction, one would presume that the carbonyl center is oriented towards the active site iron, probably in coordination with the metal ion during the course of the enzyme-catalyzed reaction. The hydrophobic pocket may assist in the binding and orientation of the substrate, while the charged residues help in directing the carbonyl group toward the Fe^+3^.

Interestingly, phenylalanine (F693) from the adjacent large subunit is part of the substrate-binding pocket. This side chain also acts as constriction and provides an explanation for selective substrate specificity and to why larger amides are not efficiently hydrolyzed (Fig. 4C). A cross-talk between the two active sites in the dimer via the loop on which F693 is present is a distinct possibility. This residue and the loop may also be involved in the observed cooperativity in the enzyme kinetic experiments. The heat of formation of the peptide bond is 2550 calories per mole and that of an amide bond is 5840 calories per mole (25). While the exact energy of breaking the amide bond in DMF has not been measured, its stability and properties that make DMF an attractive solvent would suggest that this is a very stable bond and hence its heat of formation may be even higher. When a water molecule binds to Fe^3+^, the water would make a better nucleophile that can attack the C--N bond of the formamide. Our first hypothesis was that the binding of the C=O to the Fe^3+^ will also make the carbon a better electrophile, and no other residue is involved in catalysis. However, the observation of Glu657 close to the active site suggested that it might be the catalytic base. One could think of a mechanism where the presence of the glutamate and the metal together provides the necessary catalytic power to break down the strong amide bond of DMF. We carried out site-directed mutagenesis, and the Glu657Ala enzyme did not show any activity against DMF. Together, one could propose a mechanism that involves the mononuclear Fe^3+^ center, Glu657, and the intermediate oxyanion stabilized by Asn547. Hence, the catalytic cycle will involve a water bound to Fe^3+^ as the ground state to which the substrate binds. Nucleophilic attack of the activated water on the predisposed amide bond results in the hydrolytic cleavage of the bond. This is followed by the release of dimethylamine. Formate is displaced from the active site by the binding of a new water molecule (Supplementary data Fig. 18).

While the evolutionary link to the new folds is not yet apparent, it is tempting to hypothesize that the large size of the enzyme and the properties of the quaternary structure are required to provide the enzyme with not only halostability, but is also needed for stabilization of the strained Fe^+3^ site. This strained metal coordination, the presence of the nearby glutamate that acts as the catalytic base together, provides the necessary reduction in the activation energy for increased catalytic breakdown of the substrate. Further studies to elucidate the mechanistic details of this interesting reaction are in progress. Interestingly, the use of enzymes for bioremediation has often been hindered due to lack of stability of the enzyme in the purified form or the need for other partner proteins (often to provide the necessary electronic redox potential) for catalysis. The properties of DMFase described here suggest that this enzyme might be amenable to use in bioremediation with very less genetic enzyme engineering and with no need for other proteins.

## METHODS

### Data reporting

No statistical methods were used to predetermine the sample size. The experiments were not randomized. The investigators were not blinded to allocation during the experiments and outcome assessment.

### Bacterial strains and growth conditions

Strains used in this study were either *wild-type* and/or derived from chemically competent *E. coli* One Shot™ BL21 Star™ (DE3) (F^−^ *ompT hsdS*_*B*_ *(rB*^*–*^ *mB*^*–*^*) gal dcm* (DE3)) (Invitrogen™). *wild-type(w.t.)* strain *Paracoccus sp.* strain DMF (<u>NCBI BioSample accession no. SAMN11175380</u>) was used to isolate and purify native DMFase. Cells were grown aerobically at 37 °C in an orbital shaker incubator (200 rpm) for 14 hours in modified minimal media. Compositions of modified minimal media are Na_2_HPO_4_.2H_2_O 2 g/L, KH_2_PO_4_ 1 g/L, K_2_SO_4_ 0.06 g/L, MgSO_4_ 0.25 g/L, CaCl_2_.2H_2_O 0.035 g/L, NaCl 0.5 g/L supplemented with DMF (1 g/L), yeast extract (0.01 g/L), and trace element solution (1 ml/L) {Na_4_EDTA.2H_2_O 60 g/L, ZnSO_4_.7H_2_O 22 g/L, MnSO_4_.H_2_O 5 g/L, CoCl_2_.6H_2_O 1.6 g/L, CuSO_4_.2H_2_O 1.6 g/L, H_3_BO_3_ 11 g/L, (NH_4_)_6_Mo_7_O_24_.4H_2_O 1.2 g/L, NiSO_4_.7H_2_O 1.2 g/L and FeSO_4_.7H_2_O 5 g/L (pH > 6 by adjusted by 10 N NaOH)}. For recombinant overexpression and purification of DMFase, *E. coli.* cells were grown in LB media at 37 °C.

### Plasmid construction

Standard recombinant DNA techniques and In-Fusion^®^ HD Cloning (Takara Bio USA, Inc.) were used for plasmid construction in this study and were verified by DNA sequencing. The Plasmid pET-21a (+) (*lacI*, ori F1, ori pBR322, T7 promoter and terminator, Amp^R^) was used to generate the *E. coli* overexpression vectors harboring DMFase (*w.t.*), DMFase (**β**2), DMFase (S395A), DMFase (Y399A), DMFase (Y440A), DMFase (H519A), and DMFase (E521A) with a C-terminal 6x-His tag for protein purification. The gene encoding for DMFase was generated by PCR amplification of *dmfase* gene using a bacterial plasmid isolated from *Paracoccus sp.* strain DMF as template DNA. Primers used for this study are listed in Supplementary Data Table 7. The spin-column purified PCR product was ligated with linearized pET21(a)+ vector at *NdeI* and *HindIII* cloning sites and transformed into One Shot™ TOP10™chemically competent *E. coli* (Invitrogen™) for positive clone screening and plasmid isolation.

### Protein expression and purification

DMFase was purified from both the *wild-type* and recombination sources. Cell pellets of *Paracoccus sp.* strain DMF were washed twice with **buffer A** (Tris-Cl 50 mM, NaCl 50 mM, dithiothreitol (DTT) 1 mM, pH 7.2 at 4 °C) and re-suspended in cell **lysis buffer B** (buffer A, NaCl 200 mM, MgCl_2_ 10 mM, lysozyme 0.3 mg/mL, DNase 0.01 mg/mL, glycerol 10% v/v) and incubated at 4 °C for a minimum of 4h before lysis by sonication. The cell extract was obtained by centrifugation of the lysate at 20000*g for 45 min at 4 °C. Purification was achieved by high-salt precipitation followed by ion-exchange chromatography. Ammonium sulfate salt was added to the cell extract culture (50-70% saturation range). The resulting precipitate was collected by centrifugation at 10,000*g for 10 min and re-dissolved in buffer A and dialyzed overnight against three exchanges of the same buffer. The dialyzed protein solution was incubated at 45 °C for 15 min on a heating block and was immediately transferred into ice, and heat-denatured aggregated protein fractions were removed by centrifugation at 12000*g for 15 min at 4 °C. Two-step anion-exchange chromatography was performed for further purification. The protein solution was injected into a HiPrep Q-FF (16/10) column (GE Healthcare Life Sciences) pre-equilibrated with buffer A. Column was washed with 10 CV buffer containing 0.2 M NaCl and protein was eluted with 20 CV of the same buffer with a linear gradient of 0.2 to 0.6 M NaCl. A flow rate of 1 mL/min was maintained throughout the purification process. The purified fractions as determined by SDS-PAGE (12%) were pooled and concentrated using a Millipore Amicon ultra 100K device. Dialysis was performed for 6h against buffer A and re-injected into the same column. Protein was eluted on a linear gradient of NaCl (0.3 to 0.6 M). This step was found to be essential for purification of the native DMFase to homogeneity. Peak fractions from the anion-exchange column were pooled, concentrated, and was followed by size exclusion chromatography (SEC) with a Superdex column (S-200, 16/60, GE Life Sciences) pre-equilibrated with buffer C (Tris-Cl 50 mM, NaCl 250 mM, DTT 1 mM, pH 7.2 at 4 °C). The DMFase was eluted with the same buffer at the flow rate of 1 mL/min. All the peak and pooled fractions were tested for enzymatic activity.

The DMFase constructs fused to a C-terminal His_6_ tag were expressed in *E. coli* BL21 star (DE3) cells. Cells were grown in LB medium supplemented with ampicillin (100 μg/ml) at 37 °C until they reached an OD_600nm_ of ∼0.6. Protein overexpression was induced by the addition of 0.5 mM IPTG and cultures were grown for 6h at 37°C. Cell pellets were pooled by centrifugation at 4000*g for 15 min, and resuspended in lysis buffer B and sonicated. The lysate was cleared by centrifugation at 14000*g for 45 min at 4°C. Further purification of the recombinant enzyme was performed using steps as described for native enzyme purification. SEC elution fractions corresponding to DMFase were either used immediately or flash frozen in liquid nitrogen with 20% glycerol and stored at −80 °C until further use.

### Protein concentration determination

Protein concentration at each purification step was estimated by the Bradford method using bovine serum albumin (BSA) as a standard(26).

### Steady-state enzyme assay

Amidohydrolase activity of DMFase towards *N*-substituted amides was determined using an alkylamine-specific colorimetric assay (Cullis and Waddington 1956). The standard enzyme assay with DMF contained 45 µL of 50 mM buffer C, 50 µL enzyme solution, and 5 µL of 3M DMF. The reaction was carried out in tightly closed microcentrifuge tubes for 30 min at 37°C. The reaction was quenched by the addition of 25 µL trichloroacetic acid (TCA) solution (15%, w/v). After centrifugation at 10000*g for 20 min, 100 µL of the reaction mixture was added to 1 mL of carbonate buffer (sodium tetraborate hexahydrate (9.53 g/L) and sodium carbonate (5.3 g/L) solution (pH 9.8)), 0.25 mL freshly prepared sodium nitroprusside (Na_2_[Fe(CN)_5_NO]) solution (1% w/v), and 0.25 mL acetaldehyde solution (l0% v/v). The dimethylamine (DMA) formed in the enzymatic reaction leads to a change in the absorbance, which was measured after 20 min at 580 nm. Spectrophotometric measurements were carried out with a blank solution in which DMF was added to the enzyme mixture after the enzyme inactivation step with TCA. The concentration of DMA produced upon enzymatic reaction was determined from a standard curve calibrated with known concentrations of DMA.

### Isothermal Titration Calorimetric (ITC) enzyme assay

To perform the real-time ITC enzyme kinetic assay, two sets of experiments were carried out. In single injection mode, the total molar enthalpy (ΔH°) was extrapolated by titrating 25 µL from a single injection of a relatively low concentration of substrate (25 mM) to a relatively high concentration (25 nM, 203.7 µL) of DMFase in the cell (Microcal ITC200, GE-Healthcare). In the multiple injection mode, enzymatic reaction was initiated by a series of injections of a relatively high concentration of DMF (100 mM, 2 µL, 20 injections) with a relatively low concentration of enzyme (25 nM) in the cell. All experiments were performed in buffer C at 37 °C in high-feedback mode with a stirring speed of 1000 rotations per minute and a filter time of 4s. A pre-injection delay of 600 s was applied in order to establish a steady baseline. A plot of heat rate *vs*. substrate concentration points was obtained in order to calculate the kinetic parameters after a baseline correction. The ΔH° value was determined in terms of the shift in thermal power that occurred from the conversion of the substrate before returning to its pre-equilibrated baseline(27). The Michaelis-Menten enzyme parameters K_M_, k_cat_, and steady-state heat (pseudo first order) rate were derived from the enzyme that was titrated with increasing concentrations of substrate. The kinetic experiments were carried out in duplicate. A non-linear regression fitting the Michaelis-Menten equation (GraphPad Prism v6.0 software) was used to calculate the kinetic parameters (K_m_ and k_cat_).

### ^1^H-NMR spectroscopy of enzyme kinetics for substrate specificity

Hydrolase activity of DMFase towards DMF substrate analogs including formamide, acetamide, *N*-methylformamide (NMF), *N*-ethylformamide (NEF), dimethylacetamide (DMAc), diethylformamide (DEF), benzamide, hexanamide, and urea were examined using real time ^1^H-NMR based kinetic assays. NMR experiments were carried out using Avance 600 MHz NMR spectrometer in a 5 mm probe at 310 K. All samples were prepared in 25 mM sodium phosphate buffer (pH 7.2) containing 250 mM NaCl. For the kinetics experiment, 10 nM DMFase was added to a reaction mix (500 µL) containing various concentrations of DMF and 1 mM α-alanine (internal standard). Stock solutions of DMF and α-alanine were prepared in a D_2_O assay buffer consisting of 20 mM phosphate buffer (4:1), 250mM NaCl, pH 7.2, at 37 °C. The rate of the reaction was obtained by calculating the rate of disappearance of the carbonyl peak –CH(O) of the substrate. The initial and final concentrations of the substrates were calculated by integrating the peak area of the carbonyl signal of amide with respect to the acetyl (-CH_3_) signal of α-alanine as an internal standard. The kinetic experiments were carried out in duplicate. A non-linear regression fitting the Michaelis-Menten equation (GraphPad Prism version 6.0 software) was used to calculate the kinetic parameters (K_M_ and *k*_*cat*_).

### Thermal shift assay

Protein unfolding and stability were determined as a function of temperature. In experimental studies, a label-free thermal shift assay was performed using Tycho NT. 6 (NanoTemper Technologies). Pre-dialyzed protein and mutant samples were diluted (∼0.5 mg/mL) in appropriate buffer conditions and run in duplicates in capillary tubes. Intrinsic fluorescence from tryptophan and tyrosine residues was recorded at 330 nm and 350 nm while heating the sample from 35°C to 95°C at a ramp rate of 30°C/min. The ratio of fluorescence (350/330 nm) and the inflection temperature (T_*i*_) were calculated using Tycho NT. 6 software, which provides T_m_ (melting point), the temperature at which 50% of the measured protein is unfolded. The measurement results are summarized in the Supplementary Data Table 1 and thermographs shown in Supplementary Data Fig. 6.

### Electron microscopy of DMFase

Initial images of DMFase data at 2-3 mg/ml in 50 mM NaCl were collected using Quantifoil holey carbon grids (R 0.6/1, Au 300 mesh) with blotting and freezing accomplished with a manual plunger in a cold room. This initial data was collected with Titan Krios at MRC LMB, Cambridge, and Falcon 2 detector in integration mode with the EPU software. Images showed that there were two populations, and both these were picked and subjected to reference-free 2D classification. Initial models of both the populations were individually generated either with EMAN2(28) or by Stochastic gradient descent within RELION(29) with C2 symmetry imposed. The refinement of the dimer population obtained in the integration mode gave a reconstruction of ∼7 Å, indicating that the protein behaved well and a high resolution structure can be obtained. All the data described here were collected from the enzyme obtained by recombinant expression. Enzyme purified from native source showed similar maps (data not shown).

Subsequently, data were collected with a Falcon 3 detector in counting mode at 1.07 Å sampling and images were exposed for 60 seconds with a total accumulated dose of ∼27 e/Å^2^ and dose fractionated into 75 frames, with each frame having a dose ∼0.3 e. The images were grouped into 25 frames, resulting in ∼1 e/frame, and Unblur (30) was used for alignment. This initial processing was performed in Relion 2.0. The summed images were then used for automated particle picking with Gautomatch with template derived from previous data collection, and CTF was estimated with Gctf(31). Particles were extracted with a box size of 320 pixels and subjected to 2D classification, 3D auto-refinement, per particle motion-correction, B-factor weighting, and refinement. Further 3D classification was used to improve the quality of the maps by removing bad particles. This resulted in a 3.4 Å map for the dimer and 3.8 Å for tetramer. We noticed that the views in the tetramer were not diverse, may be because of thinner ice. Hence, we collected data on the dimer.

During this period, we observed that the oligomeric state of the enzyme is affected by salt and subsequently we imaged DMFase in three different salt concentrations (no salt, 200 and 500 mM NaCl). All of the data presented here were collected at the National CryoEM facility in Bangalore with a Falcon 3 detector in counting mode at 1.07 Å sampling. The grids for these datasets were Au 300 mesh either R0.6/1 or 1.2/1.3 and made with Vitrobot Mark IV at 100% relative humidity and 18°C. The grids were blotted for 3.5 seconds. The datasets were processed with Relion 3.0, including the whole frame alignment and dose-weighting with Relion’s own algorithm. Particle picking and CTF estimation were performed using Gautomatch and Gctf. Particles were extracted with a box size of 320 pixels and subjected to 2D classification, 3D auto-refinement, per particle CTF refinement, B-factor weighting with Bayesian polishing and refinement, and subsequent 3D classification. The data sets of mutants Y440A and E521A were processed similarly. The local resolution of the maps was estimated using Resmap(32). Model building was performed with Coot(33), and the model was refined with phenix.real_space_refine(34). Although there are non-protein densities in the map, we modelled only one water molecule at the active site and the rest were not modelled. Details of the EM data and model quality are presented in Table 1.

### Crystallization, refinement, and model building

The purified native and recombinant DMFase were concentrated to 10 and 5 mg ml^-1^ respectively. Initial crystallization trials of the purified DMFase were carried out using the commercially available screens PEG/Ion, PEG/Ion 2, Crystal Screen, Crystal Screen 2 (Hampton Research) and Wizard Classic 1 and 2 (Rigaku Japan). Crystals of DMFase were obtained by mixing 200 nl of protein in buffer A with equal volumes of precipitant. All trials were conducted by hanging-drop vapor diffusion and incubated at 18°C. Diffracting protein crystals were obtained in a condition consisting of 0.1 M Tris pH 7.5, 0.2 M KCl, 18% w/v PEG 3350 after refinement of the screening conditions. The crystals were cryoprotected with reservoir buffer containing 10% ethylene glycol and flash-frozen in liquid nitrogen (*liq*. N_2_). Data were collected from single crystals at Advanced Light Source (ALS) Berkeley and subsequently scaled and reduced with XDS(35) and aimless(36). These DMFase crystals diffracted to approximately 3.5 Å and the EM model was used for molecular replacement in Phaser(37). However, no molecular replacement solution was found. The observation in cryoEM that two oligomeric states of DMFase exist in solution depending on the salt concentration prompted us to set up crystallization with the enzyme purified in the presence of high salt concentration. This yielded highly reproducible crystals that diffracted up to 2.8 Å (Table 2) and the structure was refined to a R-factor of 0.25 (R-free 0.28). The crystals belonged to space group P2_1_, and a molecular replacement solution using the tetramer model obtained from the EM data was obtained. There are very few interactions between the two tetrameric structures observed in the crystals, suggesting that this is not a higher-order oligomer.

Iterative cycles of model rebuilding and refinement were performed in COOT and Phenix.refine, respectively. Complete data collection and refinement statistics are presented in Table 2. The buried area dimer interface between the one αβ and the second αβ is ∼5000 Å^2^, and there are 21 hydrogen bonds and at least 17 salt bridges and a total of about 500 non-bonded interactions (as calculated with PDBsum(38)). This interface between the two dimers is mostly hydrophobic, with a buried surface area of approximately 1830 Å^2^ with less than 18 hydrogen bonds.

### Synchrotron X-ray fluorescence spectroscopy (syncXRF)

The identity of the metal present in DMFase was validated by XRF of protein crystals. Protein crystals were flash frozen in liquid N_2_, and sequentially washed four times in the cryoprotectant consisting of 9:1 v/v of mother liquor and ethylene glycol. XRF scans were taken at the PROXIMA-1 beamline at the SOLEIL synchrotron at 107K. The X-ray fluorescence emission spectrum of the sample was collected by excitation at the selenium K edge (12,664 eV). The spectrum contains X-ray emission lines characteristic of Fe. The peak at ∼7,100 eV corresponds to the K_α_ emission energy of iron.

### EPR study of DMFase

Protein samples for EPR were prepared by concentrating the protein up to 20 mM in buffer A. Samples were taken in the probe and frozen in liquid nitrogen. EPR spectra were obtained on a Bruker ELEXSYS-IIX-band (ν = 9.442 GHz) digital EPR spectrometer. Spectra processing and simulation were performed using a Bruker WIN-EPR and SimFonia software. A control experiment with buffer was also carried out.

### Salt, temperature, and solvent dependent DMFase stability and activity

To understand the effect of temperature and salt concentration on the activity and stability of DMFase, we performed steady-state enzyme assays, and thermal shift assays in various assay conditions. All experiments were performed in sodium phosphate buffer (25 mM, pH 7.2, NaCl 50 mM, DTT 1 mM) and in triplicate. For thermal shift measurements, enzyme at different salt concentrations, DMFase (600 nM) were first incubated for 24 hours at room temperature in phosphate buffer containing increasing NaCl concentrations ranging from 50-5000 mM. the inflection temperature (T_*i*_) of each sample was obtained by thermal shift assay performed on a Tycho NT.6 system (Supplementary Data Fig. 14). Next, we performed steady-state enzyme kinetics experiments to determine the relative activity of DMFase towards DMF (Supplementary Data Table 1). The measurements were carried out in 50 μL reaction mixture, containing final concentrations of enzyme (600 nM) and DMF (300 mM) at 37°C for 20 min. Enzymatic reaction was inhibited by protein denaturation by adding TCA, and further a colorimetric assay was performed to determine the DMA concentration as described above. Similarly, the effect of temperature on enzyme activity was measured by performing steady-state enzyme assay at different temperatures and measuring the relative catalytic activity with reference to the activity at 37°C (Supplementary Data Fig. 14).

To understand the solvent-based stability of DMFase, DMFase was incubated for at least 24h in different DMF-buffer concentrations at 4°C. The structural integrity of DMFase was measured by thermal unfolding profile using a Tycho 1.6. The enzymatic response of incubates was recorded by a time-independent assay at 37°C. The thermal unfolding characterization of incubates shows a gradual decay in mean T_*i*_ with an increased organic-aqueous ratio. Interestingly, DMF showed a stabilization effect (with increased T_*i*_ of 3°C) up to ∼7.5% v/v solvent content. A higher organic -- aqueous ratio leads to conformational instability by monitoring the blue shifts in the detected first inflection temperature. The loss of the native thermal unfolding profile occurs after 65% v/v solvent presence (>7M). Enzyme catalytic response showed gradual decay of activity as opposed to a sharp decline in higher solvent medium (Supplementary Data Fig. 16). Optimum relative activity measured for DMF was at 3.1% v/v. In high organic solvent medium (>40%), disintegration of the catalytic structure leads to the loss of total activity even with the remaining residual quaternary structure.

### Density functional theory calculations for strained iron in DMFase

We used a recently published method for calculating the electronic energies of the iron complex using density functional theory(39) with the B3LYP functional for singlet calculations, the UB3LYP method for doublet calculations, and the 6-311G(d) basis set. We used the 6-311G(d) basis set for all atoms, as the difference in computation time was not significant. We assumed a low spin state of iron in both octahedral and tetrahedral calculations. The Fe^3+^ charge state results in a doublet, and the Fe^2+^ charge state results in a singlet regardless of the tetrahedral or octahedral symmetry around the iron atom. Moreover, both tyrosine residues were assumed to be deprotonated, along with the glutamic acid residue. Therefore, the Fe^3+^ case results in a neutral system, whereas the Fe^2+^ case results in an anionic state. The initial coordinates were obtained from the crystal structure and the beta carbon was replaced with a methyl group to sever as a single bond between two SP^3^ carbons. For each geometry and charge state, two optimizations were performed: the first was constrained so that only the hydrogens and water atoms were allowed to move (strained state), and a second optimization was were all atoms were allowed to move (relaxed state). For the doublet calculations, where an unrestricted calculation was performed, the <S^2^> values were calculated before and after the annihilation of the first spin contaminant (Supplementary Data Table 2). The expected value of <S^2^> for the system lacking any spin contamination is 0.75 for the doublet system. Since <S^2^> after spin annihilation for both Fe^3+^ symmetries are 0.7559 and 0.7503 (Supplementary Data Table 2), which is relatively close to 0.75, we conclude that our calculations do not suffer from spin contamination. All calculations were performed using Gaussian16 (Frisch, M. J. *et al.* Gaussian16 Revision B.01. (2016)), and the geometries of the octahedral case for both optimizations are provided in Supplementary Data Fig. 9. For the calculation of the acetal intermediate of DMF, the B3LYP functional was used with the 6-311G(d) basis set. The initial coordinates of DMF, water, and acetal were obtained using the build molecule functionality of GaussView 6.0. All structures were checked for negative frequencies to ensure that a proper intermediate was found.

## Acknowledgments

CA thanks the Council of Industrial and Scientific Research, India for a junior and senior research fellowship. RG thanks partial support from IIT Kanpur for this work. KRV and RS thank the DBT B-Life grant DBT/PR12422/MED/31/287/2014 for major part of the funding. This research was supported by the Institute for Stem Cell Science and Regenerative Medicine (inStem) and National Center for Biological Sciences (TIFR) core funding. We also acknowledge the infra-structure grant for the in-house X-ray and NMR Facility at NCBS/inStem (BT/PR5081/INF/22/156/2012). Travel and access to the ESRF was supported by a grant BT/INF/22/SP22660/2017 from the Department of Biotechnology. KRV and AC acknowledge the MRC for support in the initial stage of the project. KRV thanks SERB, India for the Ramanujan Fellowship. GC acknowledges start-up funds from the Department of Chemistry and award from the Integrative Data Science Institute at Purdue University. JF acknowledges the bioinformatics fellowship from Purdue University Center for Cancer Research.

## Contributions

RG and RS conceived the project. RG, RS, and KRV designed the experiments. SY, AC, and KRV carried out sample preparation, EM data collection analysis. KRV built the initial model. CA and RS carried out the X-ray structure determination and analysis. CA carried out most of the biochemical studies. JF and GC carried out DFT calculations. CA, RS, RG, GC, and KRV wrote the manuscript. All authors reviewed and corrected the manuscript.

