## Supplementary Data for "Breaking a Strong Amide Bond: Structure and Properties of Dimethylformamidase" (from another version of this preprint: version 1, 2019-12-18)

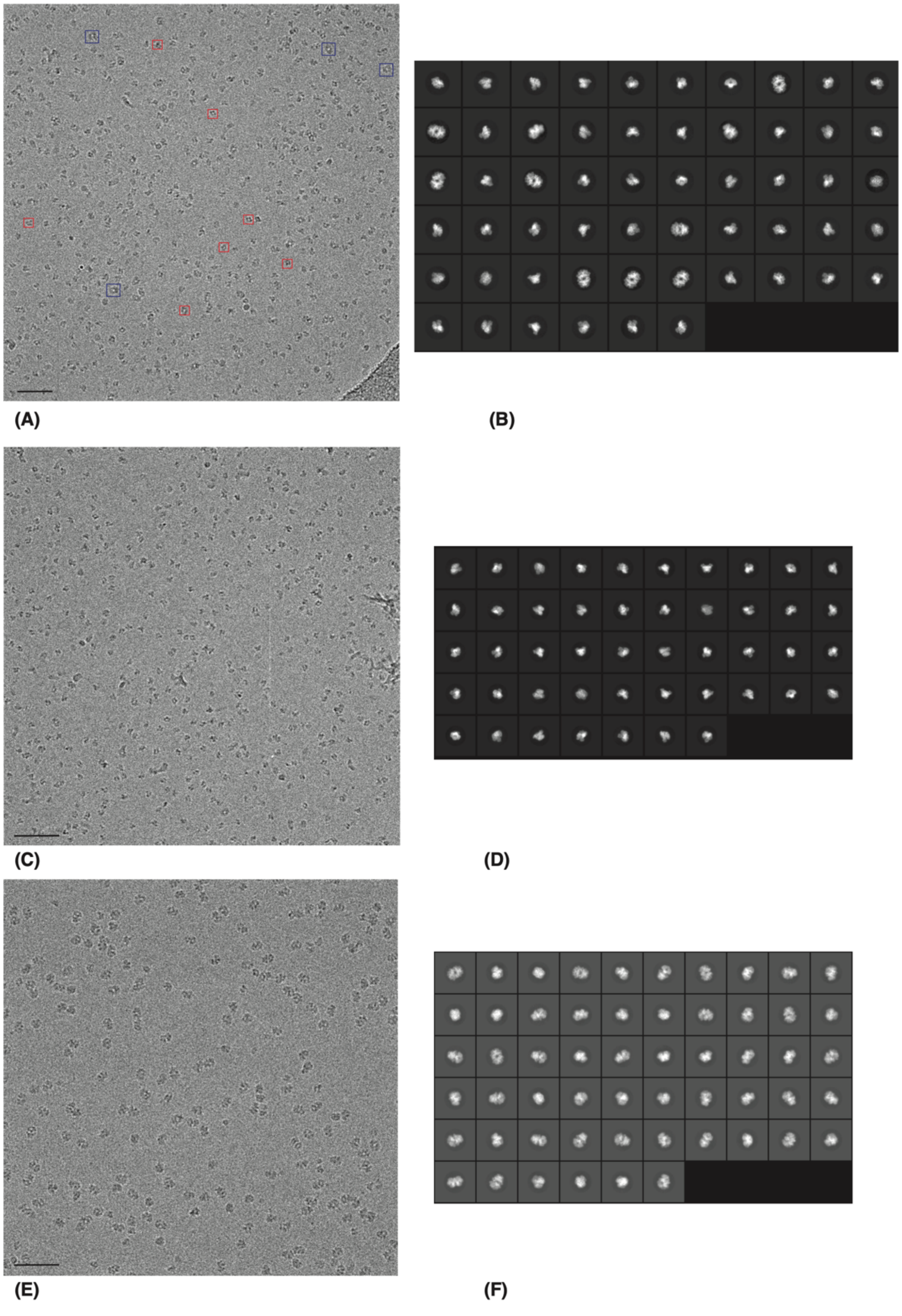
(A)
(B)
(C)
(D)
(E)
(F)

**Figure 1: CryoEM micrographs and 2D classifications**

(A) Micrograph of DMFase that shows mixed population. The smaller molecule (shown in red box) and larger molecule (blue box) are observed. Scale bar is 500 Å.

(B) All the particles from mixed populations was picked and subjected to reference-free 2D classification, both the smaller and large molecule could be separated. Box size is 256 pixels and sampled at 1.38 Å/pixel.

(C) DMFase imaged with no salt in the buffer. This shows predominantly dimer population. Scale bar is 500 Å.

(D) Representative reference-free 2D class averages of the molecule with no salt buffer. Box size is 320 pixels and sampled at 1.07 Å/pixel.

(E) DMFase imaged with 200 mM of NaCl in the buffer. This shows predominantly tetramer population. Scale bar is 500 Å.

(F) Representative reference-free 2D class averages of the molecule with no salt buffer. Box size is 320 pixels and sampled at 1.07 Å/pixel.

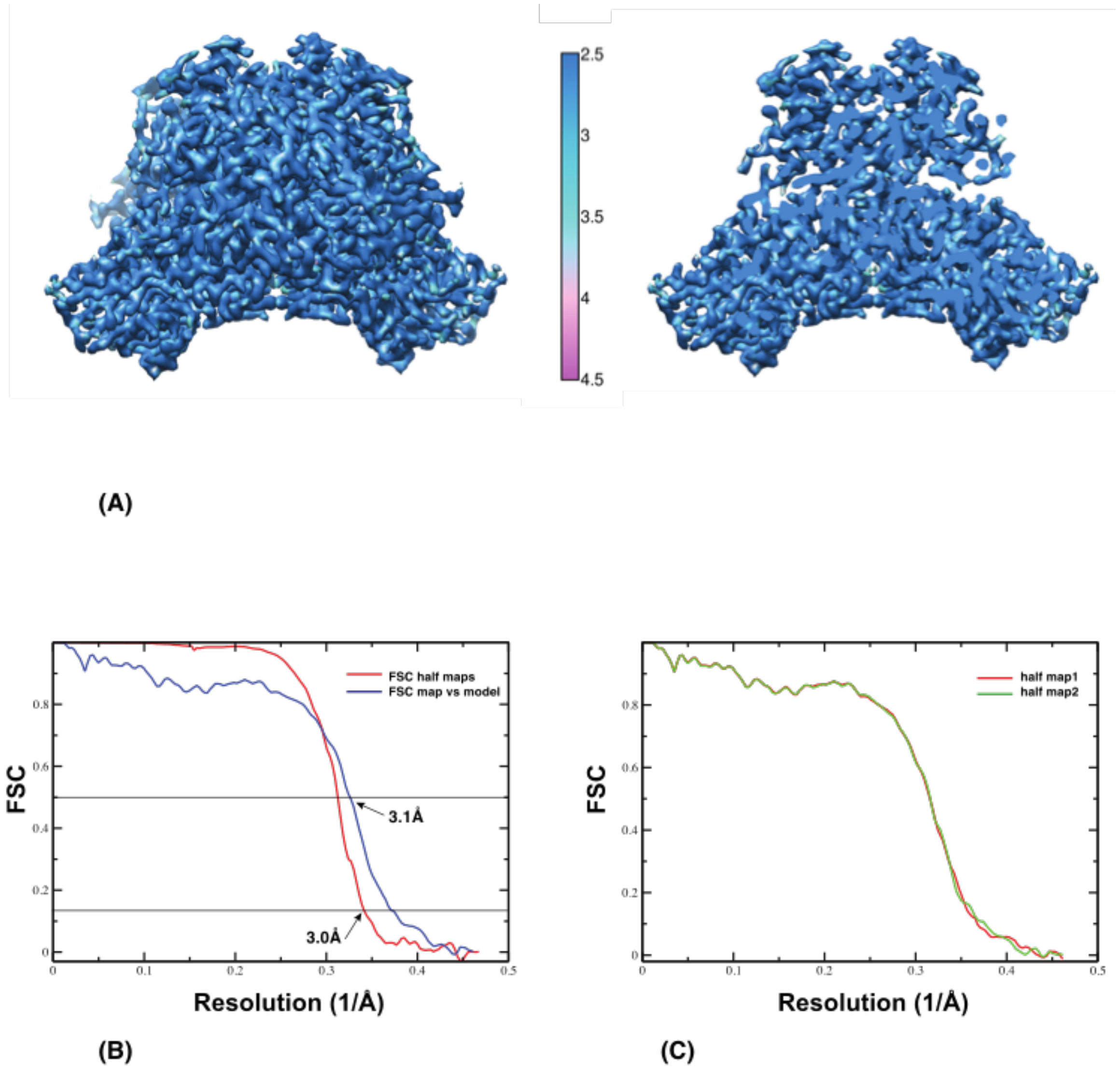


**Figure 2: Local resolution of the DMFase dimer and the FSC plots**

(A) The local resolution plot of DMFase dimer showing the whole molecule on right and a cut away section on left. Much of the molecule shows resolution between 2.5-3.5 Å allowing the building of the complete model.

(B) The Fourier shell correlation (FSC) curves of two half-maps (red) and the map and model (blue) are shown.

(C) To verify overfitting of the model refinement, one of the half-map used in refinement and the other half-map was used as test.

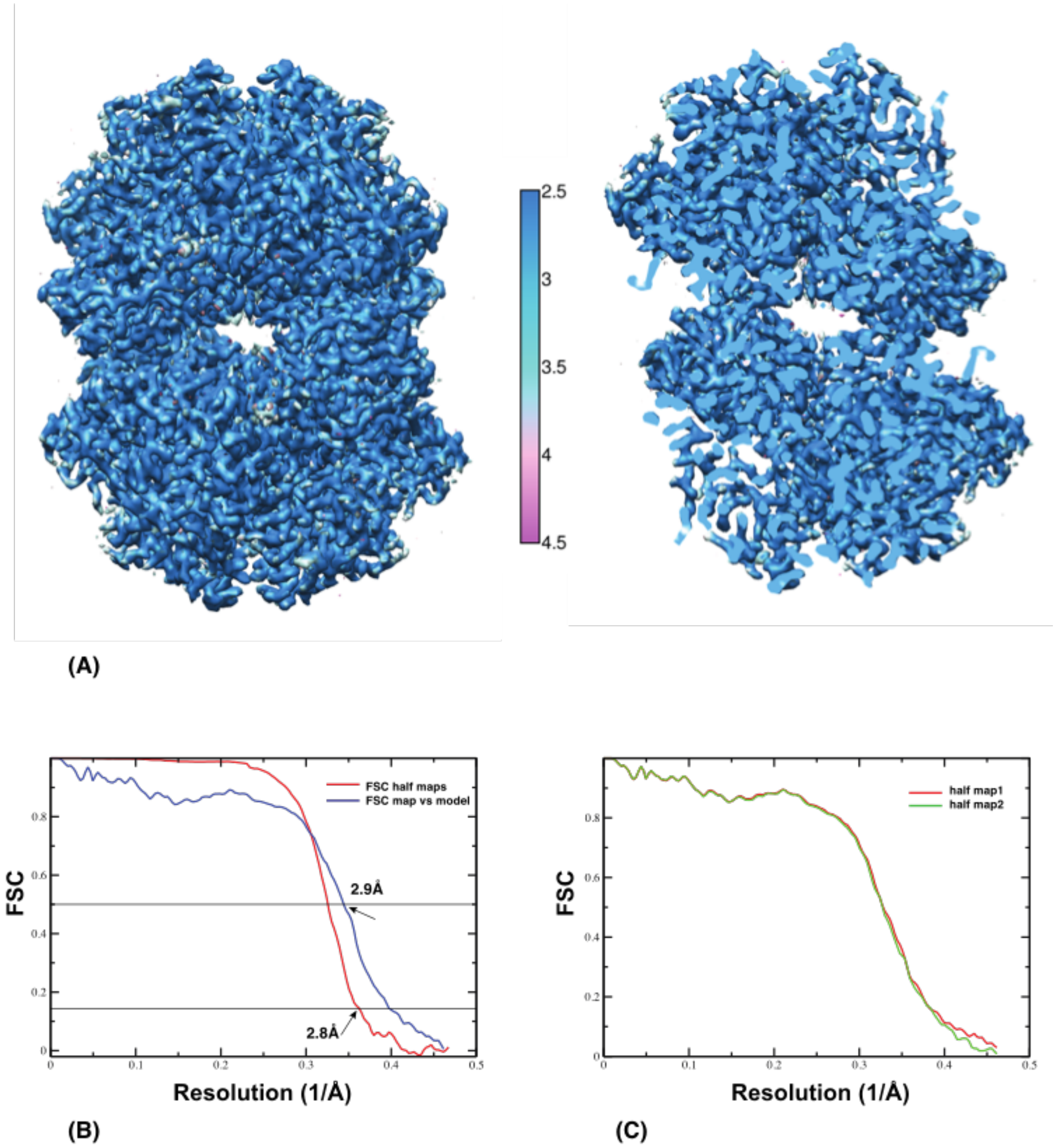


**Figure 3: Local resolution of the DMFase tetramer and the FSC plots**

(A)The local resolution plot of DMFase tetramer showing the whole molecule on right and a cut away section on left. Much of the molecule shows resolution between 2.5-3.5 Å allowing the building of the complete model.

(B)The Fourier shell correlation (FSC) curves of two half-maps (red) and the map and model (blue) are shown.

(C) To verify overfitting of the model refinement, one of the half-map used in refinement and the other half-map was used as test.

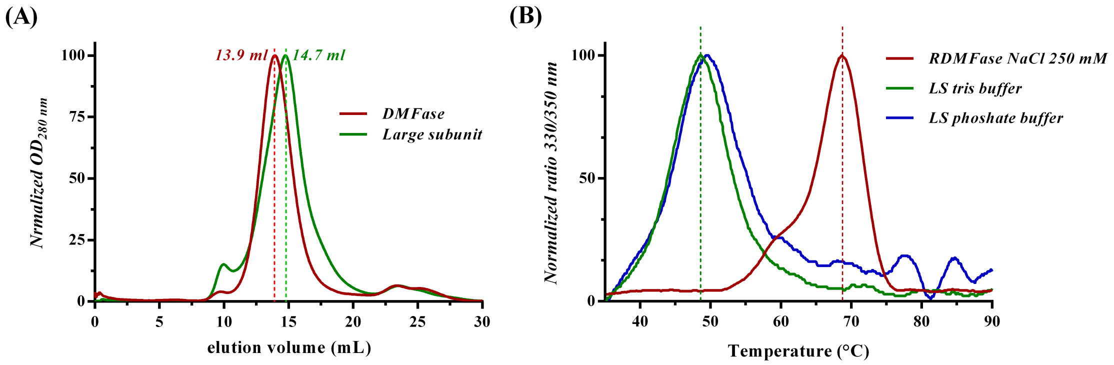


**Figure 4: Large subunit SEC profile and inversion temperatures; (A)** Size exclusion profile of recombinant DMFase(RDMFase) dimer (**in red**) and large subunit (LS) alone (**in green or blue**). Calculated molecular weights of DMFase and large subunit correspond to 200 and 172 kDa and the elution profile indicate that large subunit alone also exists in dimer. **(B)** Inversion temperatures of purified large subunit in different buffers (tris-NaCl and sodium phosphate buffer with NaCl 250 mM) and comparison with DMFase. A significant decrement in $T_i$ suggests a less stable oligomeric state of the large subunit alone than the $\alpha_2\beta_2$ dimer.

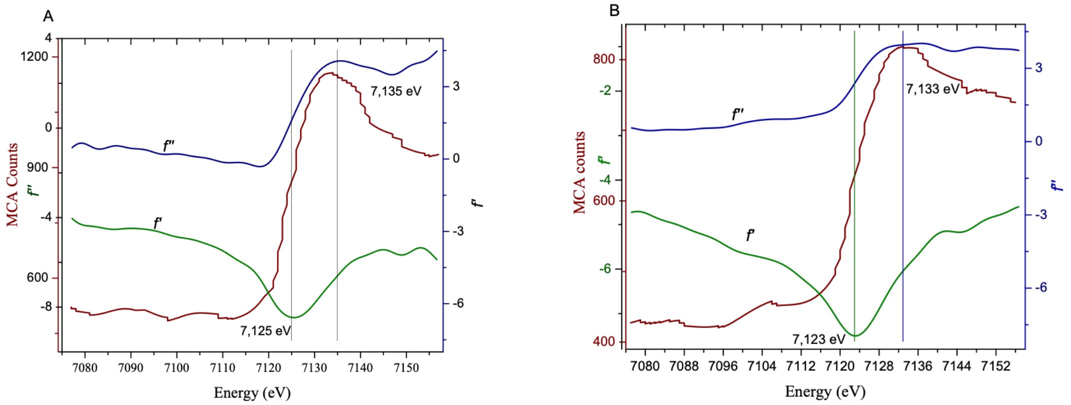


**Figure 5**: X-ray fluorescence scan of (A) native and (B) recombinant DMFase crystals at the Iron K-edge region. The *f″* plot and the observed absorption edge (7,135 eV or 2.10 Å) correspond to values for the iron $K_{\alpha}$ edge.

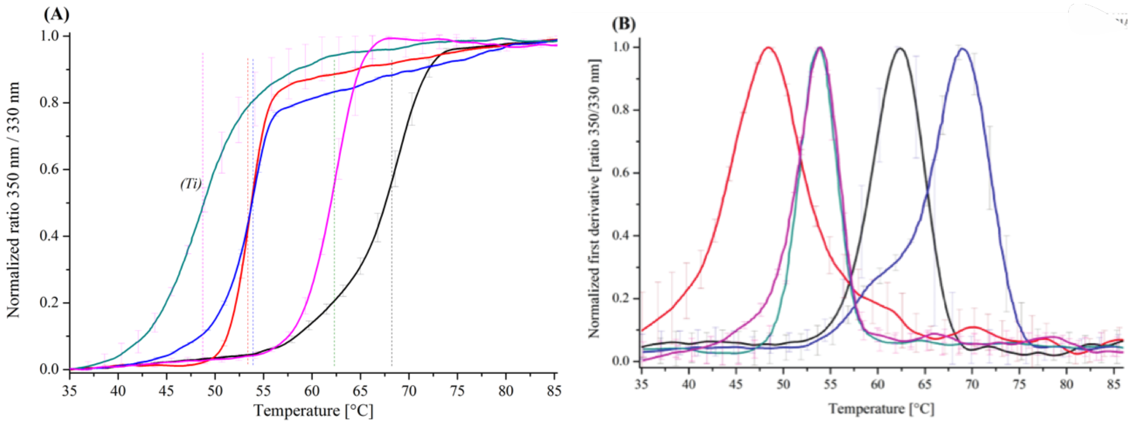


**Figure 6.** Unfolding profiles of *wild-type* and mutants of DMFase in Buffer C. (**A**)**.** Normalized thermograph of ratio of the 350 nm/330 nm fluorescence signal recorded against temperature gradient. (---) *wild-type* protein; (---) S395A; (---) Y440A; (---) H519A; (---) E521A; (**B**)**.** First derivative graph of (A).

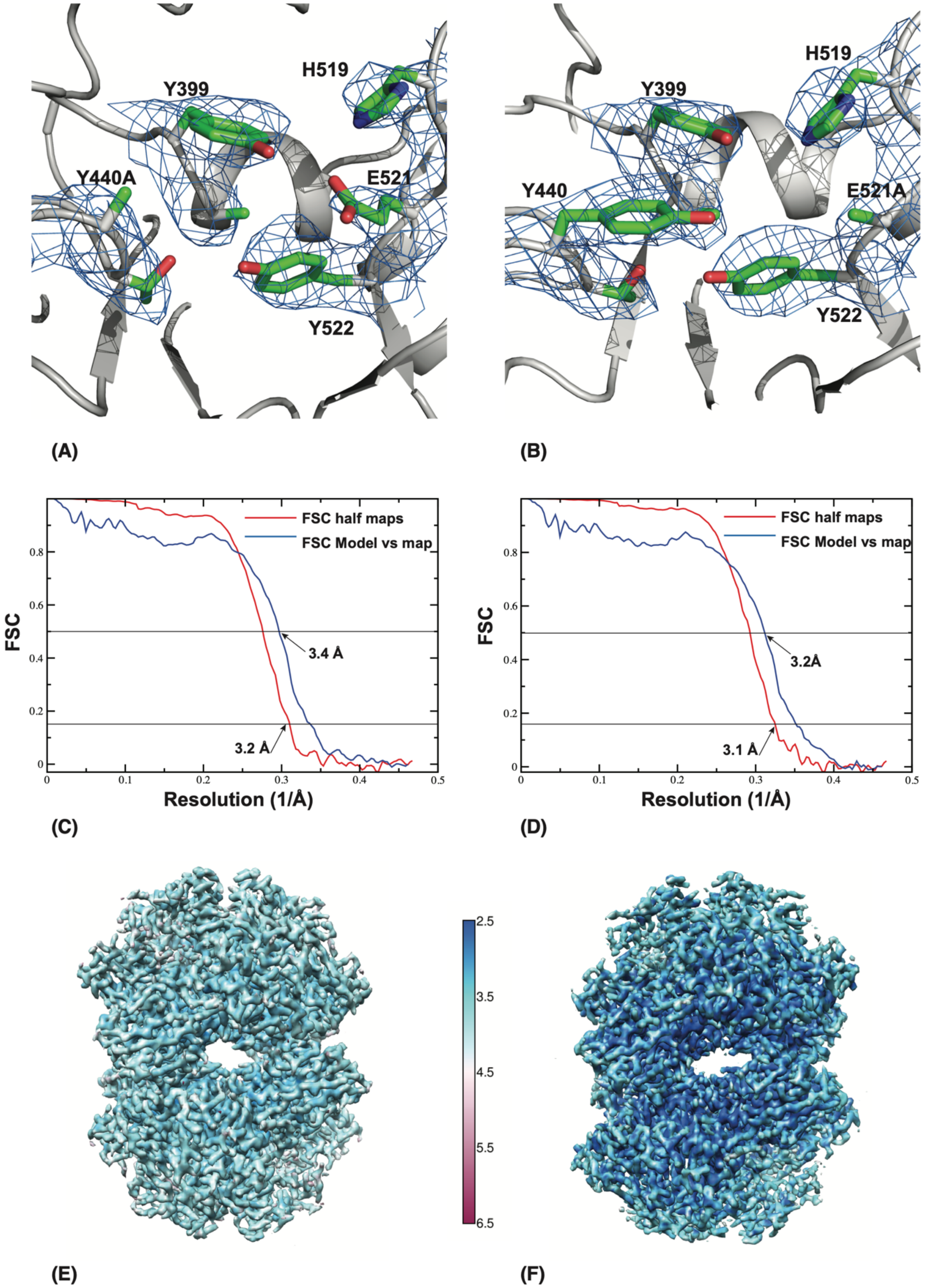
H519
Y399
Y440A
E521
Y522
(A)
H519
Y399
Y440
E521A
Y522
(B)
FSC half maps
FSC Model vs map
FSC
0.8
0.6
0.4
0.2
0
3.4 Å
3.2 Å
0
0.1
0.2
0.3
0.4
0.5
Resolution (1/Å)
(C)
FSC half maps
FSC Model vs map
FSC
0.8
0.6
0.4
0.2
0
3.2Å
3.1 Å
0
0.1
0.2
0.3
0.4
0.5
Resolution (1/Å)
(D)
2.5
3.5
4.5
5.5
6.5
(E)
(F)

**Figure 7:** Cryo-EM structures of Y440A and E521A

(A & B) The active site of the Y440A and E521A mutants. Density around the active site residues that coordinate Fe are shown. In both these mutants, the extra density that is seen in wild type enzyme is absent.

(C & D) The Fourier shell correlation (FSC) curves of two half-maps (red) and the map and model (blue) are shown for DMFase Y440A and E521A mutants respectively.

(E & F)The local resolution plots of DMFase Y440A and E521A mutant respectively.

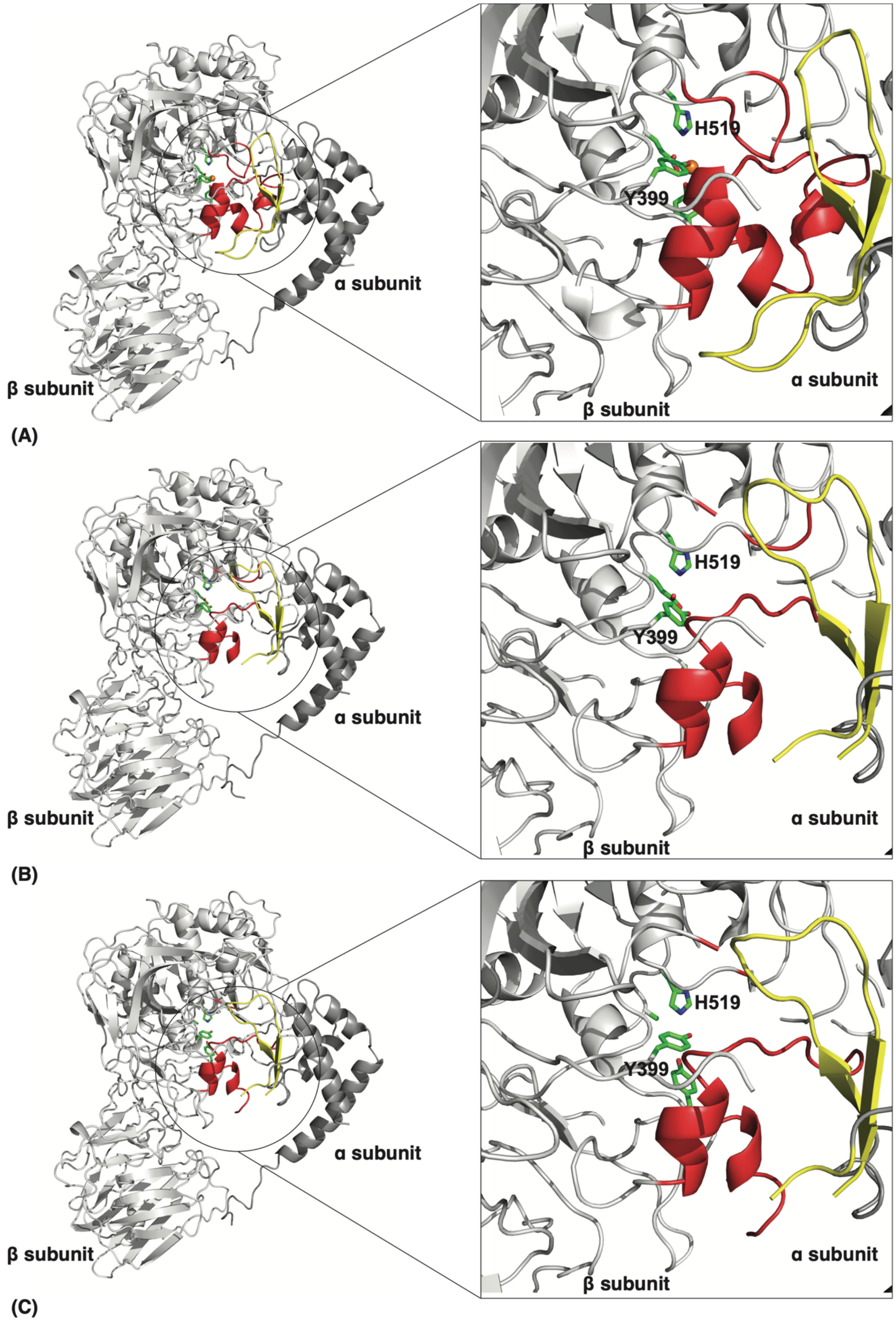


**Figure 8**: The interface of large and small subunit in wild type and Y440A, E521A mutants. On right, a monomer of DMFase is shown in different shades of grey colour for the large and small subunits. The interface region between the large and small

subunit is shown in red and yellow respectively. On the left, a zoomed view of the interface region is shown for all the enzymes wild-type (panel A), Y440A (panel B) and E521A (panel C). The active site residues Y399, Y440, E521 and H519 (stick representation) are shown along with iron molecule (sphere) in all three panels. Residues corresponding to 410-415 and 467-470 in the large subunit and 96-100 in the small subunit become disordered in the mutant enzymes.

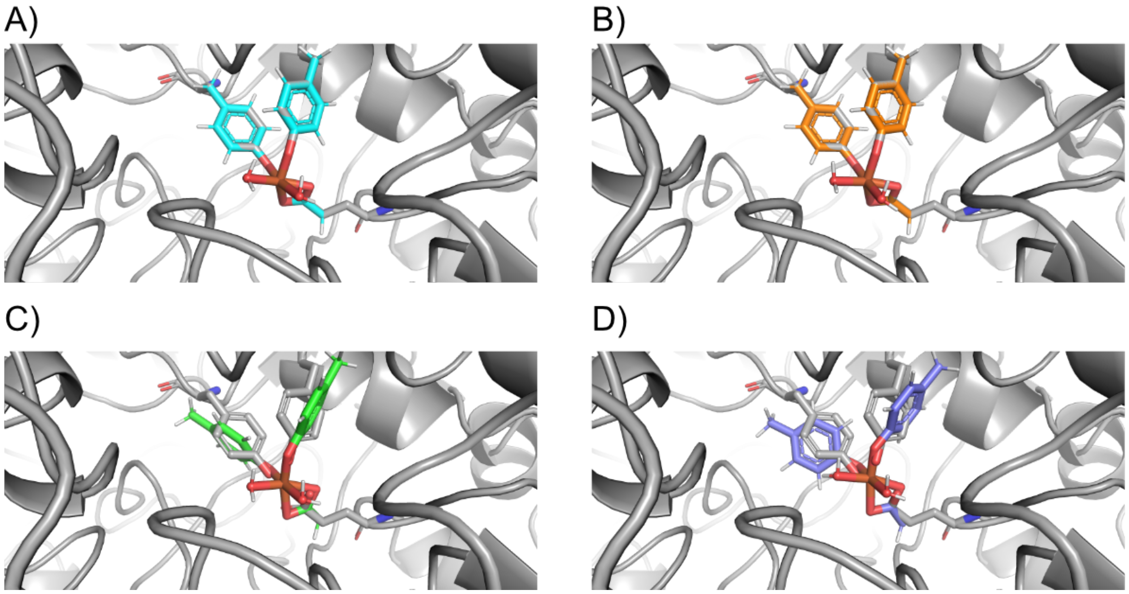


**Figure 9:** The strained conformations of the binding site iron in an octahedral conformation with charges of 2 and 3 on the iron atom are given in (A) and (B), respectively. The relaxed conformations of these $Fe^{2+}$ and $Fe^{3+}$ complexes are given in (C) and (D), respectively.

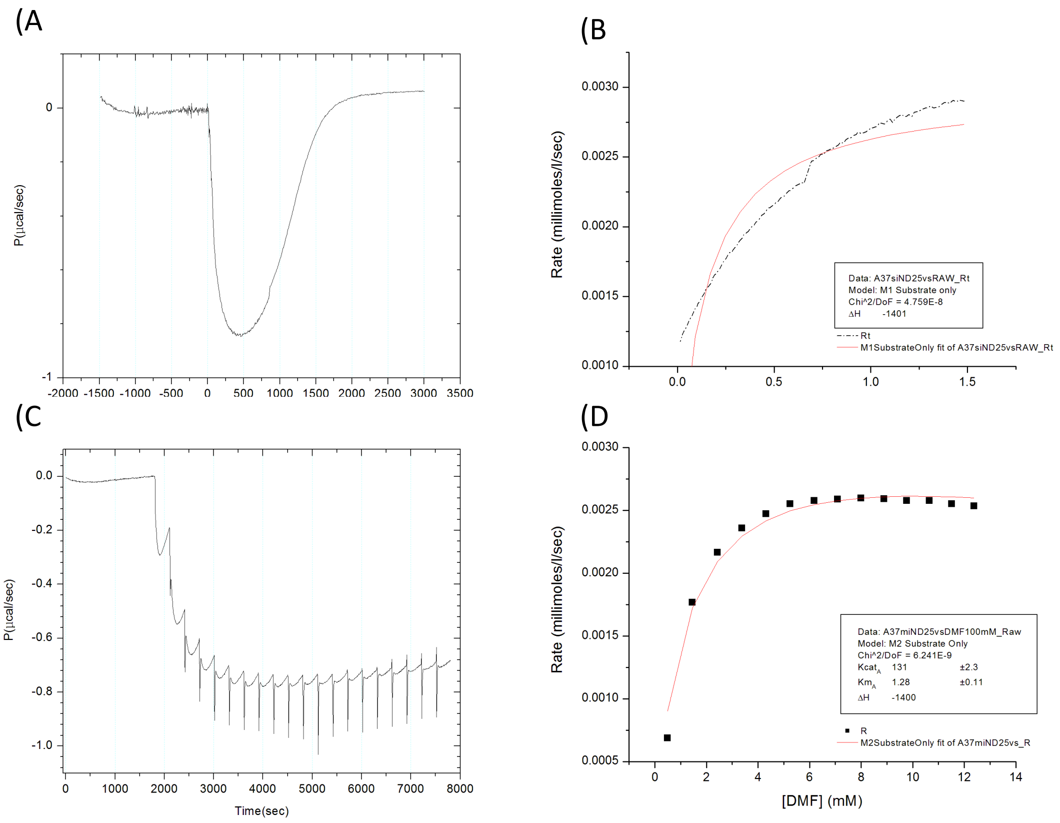


**Figure 10:** ITC kinetic assay for native DMFase catalyzed hydrolysis. (A) Raw data for thermal power (µcal/sec) single injection thermographs at 37ºC. (B) Data from (A) has been transformed into reaction rates vs. substrate concentration [S] (mM) with the nonlinear least squares fit to the Michaelis-Menten equation for ΔH determination. Calorimetric assay thermograph; (C) Raw data from Multiple injection method for thermal power (µcal/sec) at 37ºC. (D) $K_m$ and $k_{cat}$ with the non-linear least squares fit to the Hill equation from plot rate vs. [DMF].

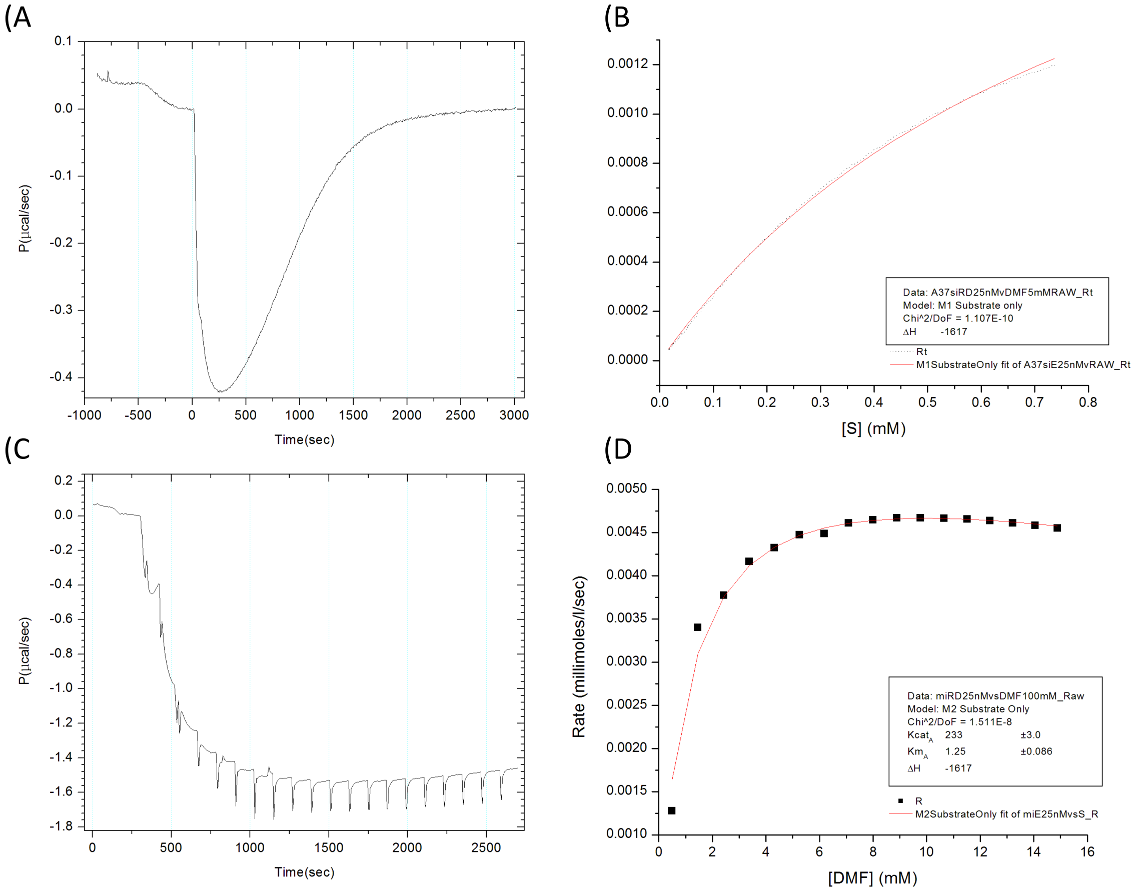


**Figure 11:** Normalized ITC single injection thermographs for recombinant DMFase catalyzed hydrolysis. (A) Raw data for thermal power (µcal/sec) at 37ºC. (B) Data from (A) has been transformed into reaction rates (millimoles/I/sec) vs. substrate with the nonlinear least squares fit to the Michaelis–Menten equation for ΔH determination. Calorimetric assay thermograph; (C) Raw data from Multiple injection method for thermal power (µcal/sec) at 37ºC. (D) $K_M$ and $k_{cat}$ with the non-linear least squares fit to the Hill equation from plot rate vs. [DMF].

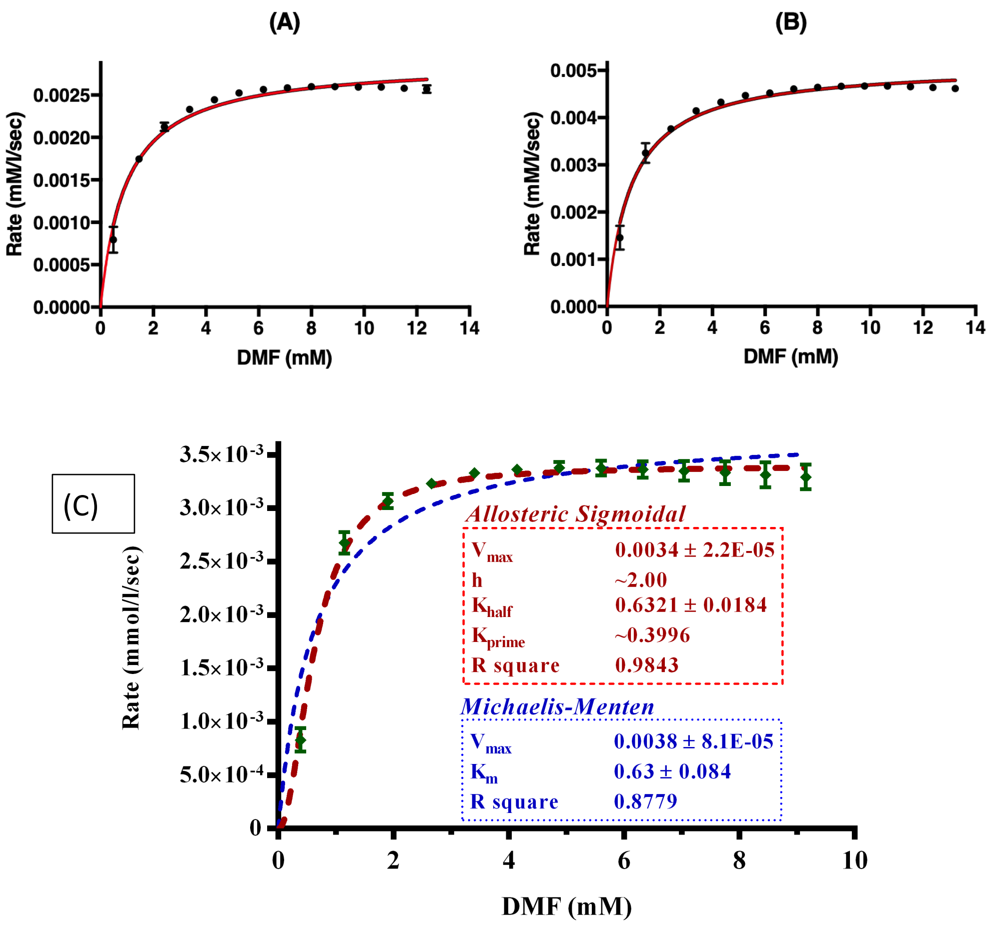


**Figure 12:**

Real-time ITC kinetic assay of (A) native and (B) recombinant DMFase. All data were fitted to the Michaelis-Menten equation $V_0 = V_{max}[S]/K_m + [S]$ using GraphPad Prism v6.0. All fits to Michaelis-Menten equation resulted in $r^2 > 0.97$. (C) Evidence of intra-molecular cooperativity between two active sites within the dimer with Hill's coefficient (h = 2).

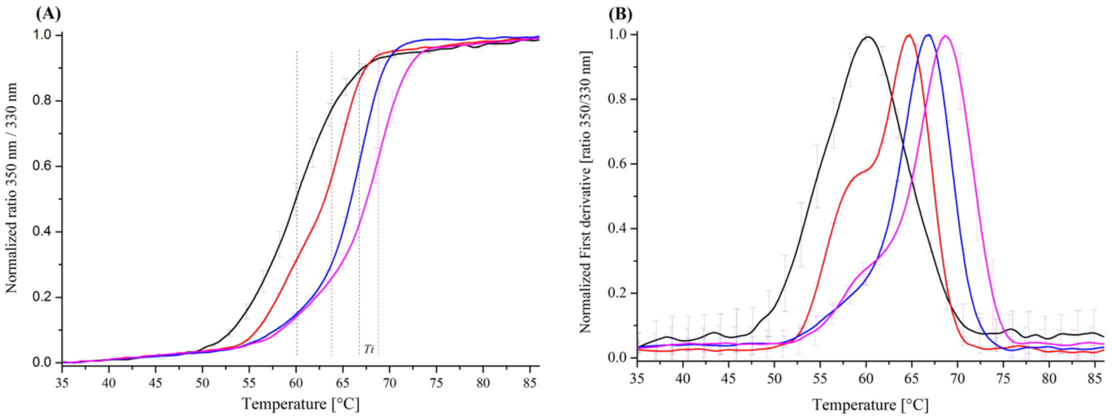


**Figure 13.** Unfolding profiles of DMFase in different buffer-salt conditions. (**A**). Normalized thermograph of ratio of the 350 nm/330 nm fluorescence signal recorded with respect to temperature gradient. (---) protein in tris buffer pH 7.2, 50 mM NaCl; (---) tris buffer pH 7.2, 250 mM NaCl; (---) tris buffer pH 7.2, 500 mM NaCl; (---) phosphate buffer pH 7.2, 250 mM NaCl; (**B**). First derivative analysis of (A).

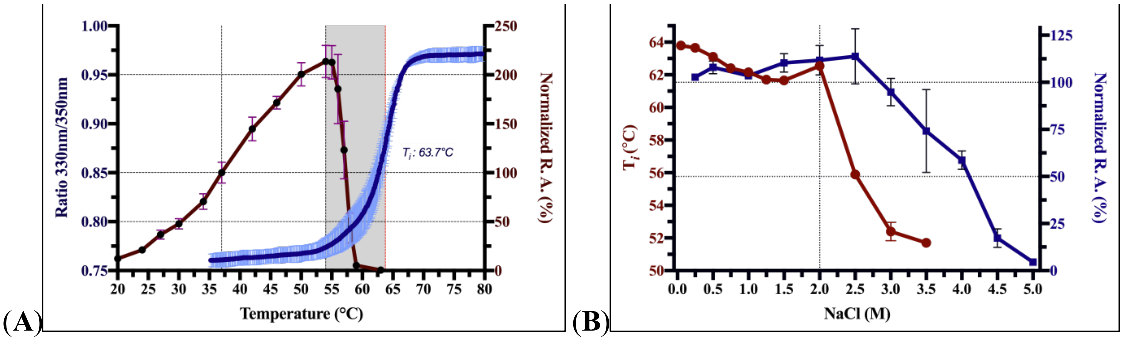


**Figure 14:** Hydrolytic activity of DMFase at different temperatures and NaCl concentrations. All assay solutions (50 µl) contained 300 mM DMF and 0.6 mM DMFase in sodium phosphate buffer supplemented with 250 mM NaCl. The assay was performed for 20 min. **(A)** Relative catalytic activity (R.A) (---) was measured with respect to amidase activity at 37°C. The raw thermal unfolding of DMFase (---) plotted of intrinsic fluorescence response with increased temperature. **(B)** The $T_i$ of the enzymes incubated in different salt concentrations plotted (---) with respective relative amidase activities (---).

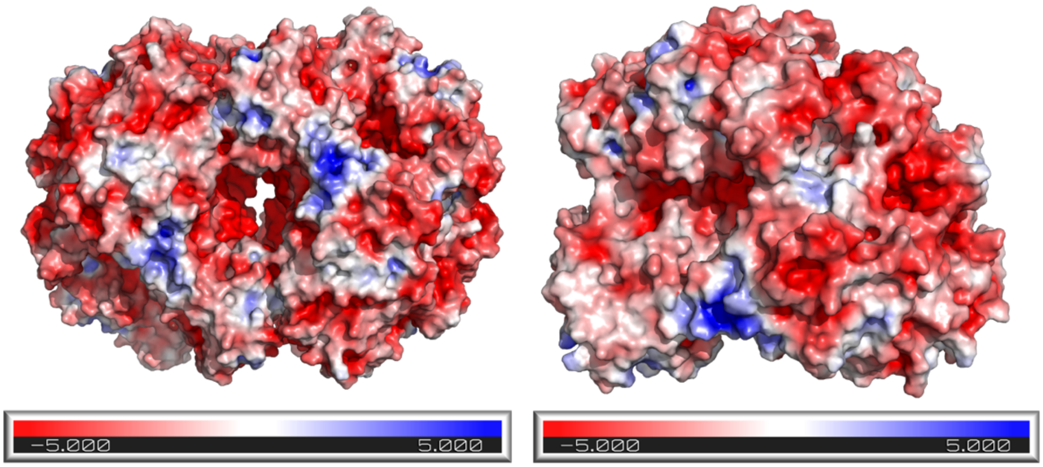


**Figure 15:** Surface charge distribution of tetramer and dimer form of DMFase. The electrostatic potential is represented in solid surface; the isosurfaces were set in +5 mV (blue) and -5 mV (red).

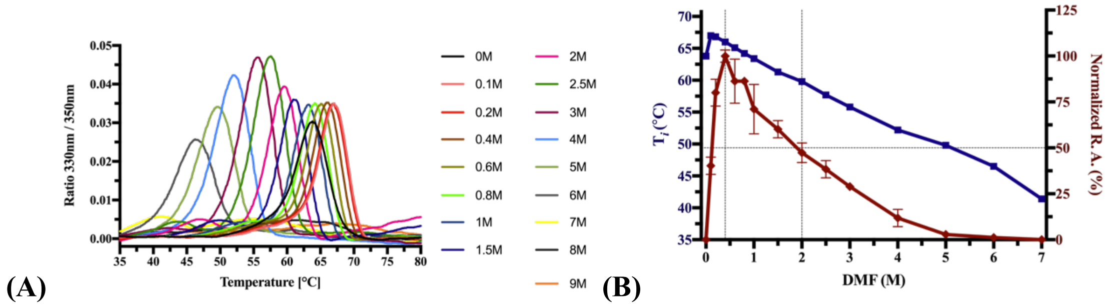


**Figure 16:**

Effect of DMF as an organic solvent on the catalytic activity and stability of DMFase. All assay solutions carried in sodium phosphate buffer (NaCl 50 mM) 0.6 mM DMFase in Na-phosphate buffer. The discontinuous assay was performed at 37°C for 20 min. **(A)** Inversion temperature (T*i*) measurement of DMFase in different concentrations of DMF in sodium phosphate buffer (50 mM, pH7.2, NaCl 50mM). Graph plotted between intrinsic fluorescence of protein and protein unfolding with increase in temperature. Blue-shift in Ti was quantifiable up to 7 M (~65% v/v) DMF concentration. **(B)** The $T_i$ of the enzymes incubated in different DMF concentrations plotted (---) with respective relative amidase activities (---).

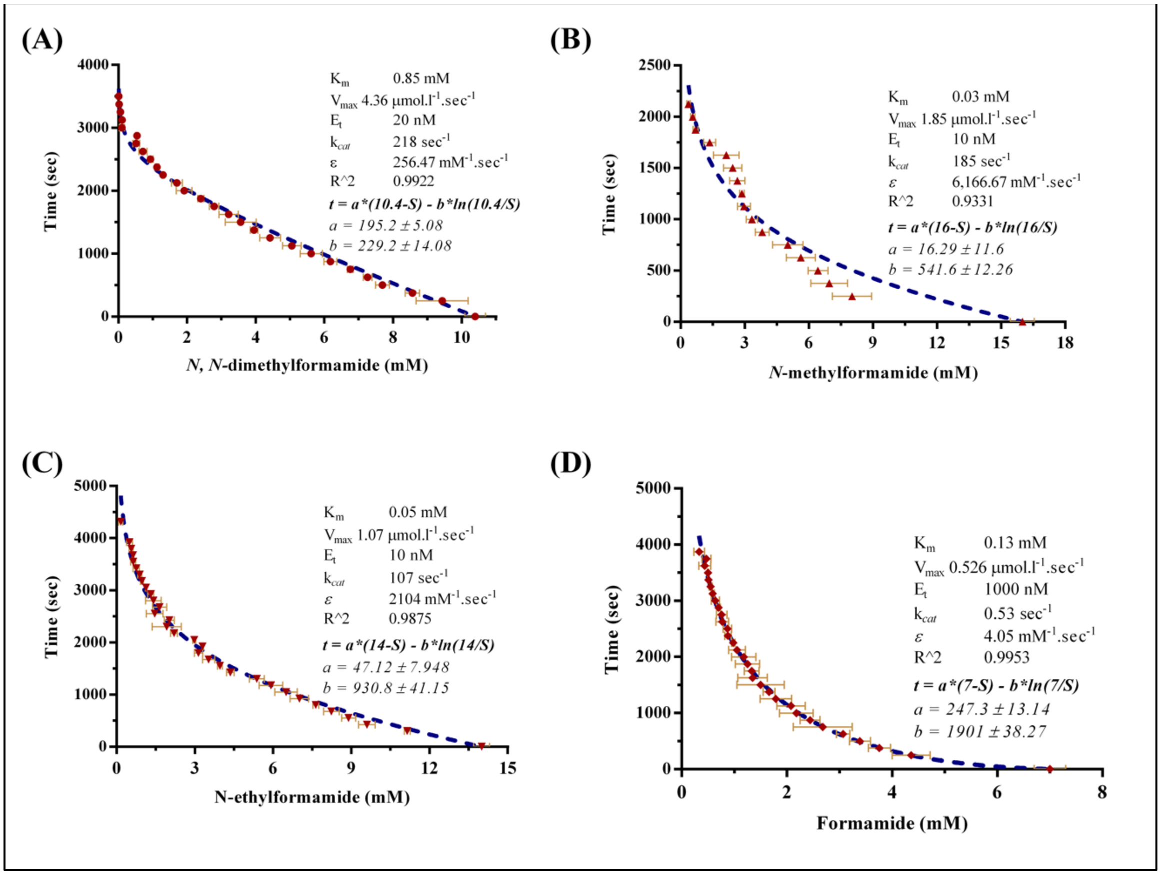


**Figure 17**: $^{1}$HNMR spectroscopy for real time kinetics for different substrates analogous to DMF. All experiments were performed at 37°C with alanine (1 mM) as internal standard in sodium phosphate buffer supplemented with 50 mM NaCl at pH 7.2. Estimation of kinetic parameters ($V_{max}$ and $K_m$) were obtained through integrated Michaelis Menten equation {$t = (K_m/V_{max})(S_0\text{-}S) - (1/V_{max}).ln(S_0/S)$: where t: time in sec; $S_0$: initial substrate concentration (mM)}. Residual substrate concentration (S) were calculated by area under curve with respect to peak corresponds to carbonyl proton.

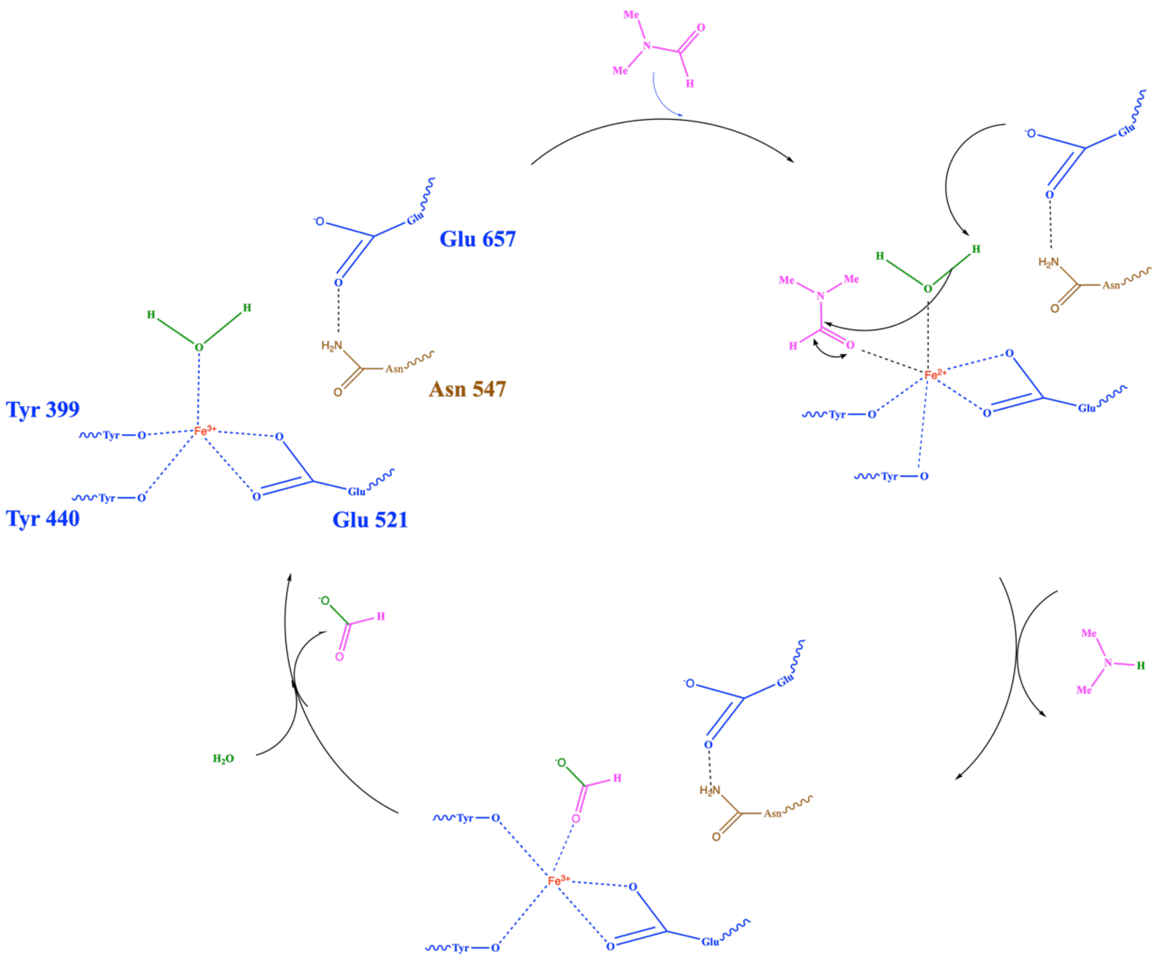


**Figure 18**: Proposed catalytic steps in the hydrolytic cleavage of DMF by DMFase. The resting state of the enzyme is the water bound to Fe+3. This is followed by the binding of the substrate to the metal center. The activated water acts as a nucleophile and attacks the C-N bond resulting in product formation. This is followed by release of the dimethylamine. A water then displaces the formate and the cycle is completed.

**Table 1. Inversion temperature and relative activity of large subunit and mutants of DMFase**

| Sample | Condition | $T_i$ (°C) [a] | Amidase Activity |
|---|---|---|---|
| **DMFase** | NaCl 50 mM, buffer C, pH 7.2 | 60.2 | + |
| **DMFase** | NaCl 250 mM, buffer C, pH 7.2 | 68.7 | + |
| **DMFase** | NaCl 500 mM, buffer C, pH 7.2 | 64.5 | + |
| **DMFase** | NaCl 250 mM, Phosphate buffer, pH 7.2 | 66.8 | + |
| **S395A** | Buffer C | 62.4 | + |
| **H519A** | Buffer C | 48.5 | _ |
| **Y440A** | Buffer C | 53.4 | _ |
| **E521A** | Buffer C | 53.6 | _ |
| **Large subunit** | NaCl 500 mM, buffer C, pH 7.2 | 46.4 | _ |

[a] $T_i$, inflection temperature obtained by automated data analysis by Tycho NT. 6 at pH 7.2. The values represent the mean ± s.d. of duplicates.

**Table 2.** Electronic energies of the iron's electronic environment. All values in atomic units unless otherwise noted. The <$S^2$> values before and after the annihilation of the first spin contaminate are separate with a '/'.

| Optimization | $Fe^{3+}$ Tetrahedral (Doublet) | $Fe^{3+}$ Octahedral (Doublet) | $Fe^{2+}$ Tetrahedral (Singlet) | $Fe^{2+}$ Octahedral (Singlet) |
|---|---|---|---|---|
| **Hydrogens and water flexible, strained state (Hartree)** | -2261.2005 | -2337.6695 | -2261.2406 | -2337.7092 |
| **All atoms flexible, relaxed state (Hartree)** | -2261.2843 | -2337.7540 | -2261.3356 | -2337.8059 |
| **Difference (Hartree)** | -0.083848 | -0.084562 | -0.095001 | -0.096712 |
| **Difference (kJ/mol)** | -220.1429 | -222.0175 | -249.4251 | -253.9174 |
| **Spin contamination** | 1.0345 / 0.7559 | 0.7811 / 0.7503 | N/A | N/A |

**Table 3**. **Real time kinetic parameters of DMFase at 37ºC**

| Enzyme | DMFase (nM) | DMF (mM) | ΔH (kcal/mol) | $k_{cat}$ ($s^{-1}$) | $K_m$ (mM) | $V_{max}$ | $k_{cat}/K_m$ |
|---|---|---|---|---|---|---|---|
| Native DMFase | 25 | 100 | -1.4 | 115 (±1.5$e^{-6}$) | 0.97 (±0.06) | 2.89 | 118 |
| Recombinant DMFase | 25 | 100 | -1.6 | 202 (±2.0$e^{-6}$) | 0.87 (±0.07) | 5.13 | 232 |

**Table 4. Relative activity and inversion temperature of DMFase in Different NaCl concentrations**

| No. | NaCl (mM) | Relative Activity (%) | $T_i$ (°C ±0.1) |
|---|---|---|---|
| 1 | **250** | **100 ± 7** | **64.0** |
| 2 | **500** | **102 ± 10** | **63.1** |
| 4 | **1000** | **102 ± 1** | **62.1** |
| 5 | **1500** | **108 ± 5** | **61.8** |
| 6 | **2000** | **104 ± 13** | **62.5** |
| 7 | **2500** | **110 ± 11** | **56.0** |
| 8 | **3000** | **94 ± 5** | **52.3** |
| 9 | **3500** | **74 ± 12** | **51.7** |
| 10 | **4000** | **57 ± 4** | **N.A.** |
| 11 | **4500** | **18 ± 4** | **N.A.** |
| 12 | **5000** | **4 ± 1** | **N.A.** |

**Table 5**. **Relative activity and inversion temperature of DMFase in Different DMF concentrations**

| **No.** | **[DMF]**<br>**mM / % v/v** | **Relative Activity [A]**<br>**(%)** | **$T_i$ (ºC)[b]** |
|---|---|---|---|
| **1** | 0 / 0 | 0 | 63.8 |
| **2** | 100 / 0.77 | 40 ± 4 | 67.0 |
| **3** | 200 / 1.5 | 80 ± 6 | 66.8 |
| **4** | 400 / 3.1 | 1.0 ± 3 | 66.0 |
| **5** | 600 / 4.6 | 87 ± 8 | 65.1 |
| **6** | 800 / 6.2 | 86 ± 1 | 64.2 |
| **7** | 1000 / 7.7 | 71 ± 1 | 63.4 |
| **8** | 1500 / 11.4 | 60 ±4 | 61.3 |
| **9** | 2000 / 15.4 | 47 ± 4 | 59.8 |
| **10** | 2500 / 19.3 | 38 ± 4 | 57.7 |
| **11** | 3000 / 23.1 | 29 ± 2 | 55.8 |
| **12** | 4000 / 30.8 | 12 ± 4 | 52.2 |
| **13** | 5000 / 38.7 | 3 ± 1 | 49.8 |
| **14** | 6000 / 46.4 | 1 ± 0 | 46.5 |
| **15** | 7000 / 54.2 | 1 ± 0 | 41.1 |

[*A*] Normalized relative activity against highest activity measured at 400 mM DMF concentration at 37 ºC in sodium phosphate buffer (50 mM, pH 7.2, NaCl 50 mM, DTT 1 mM).
[*B*] DMFase was incubated in DMF at 4ºC ( no enzyme activity was observed) overnight in previous mentioned buffer.

**Table 6. Hydrolase activity of DMFase for different analogous amides**

| Substrate | Structure | [a]$V_{max}$ | [b]$K_m$ | [c]$k_{cat}$ | [e] Catalytic efficiency |
|---|---|---|---|---|---|
| DMF | 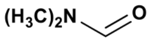 | 4.36 | 0.85 | 218 | 256.5 |
| Formamide | 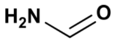 | 0.526 | 0.13 | 0.53 | 4.05 |
| NMF | 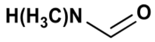 | 1.85 | 0.03 | 185 | 6166.7 |
| NEF | 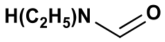 | 1.07 | 0.05 | 107 | 2104 |
| DMAc | 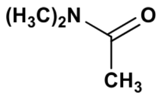 | 0.3 | *N.A.* | *N.A.* | *N.A.* |
| Benzamide | 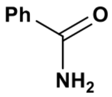 | 0 | [d]*N.A.* | *N.A.* | *N.A.* |
| Acrylamide | 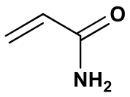 | 0 | *N.A.* | *N.A.* | *N.A.* |
| Hexenamide | 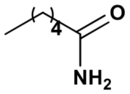 | 0 | *N.A.* | *N.A.* | *N.A.* |
| Urea | 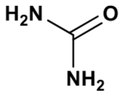 | 0 | *N.A.* | *N.A.* | *N.A.* |
| Acetamide | 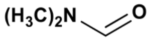 | 0.2 | *N.A.* | *N.A.* | *N.A.* |

***a)***: $\mu mol/l^{-1}.sec^{-1}$; ***b)***: mM; ***c)***: $sec^{-1}$; ***d):*** not available; ***e)*:** $mM^{-1}.sec^{-1}$

**Table 7**. **Primers used in this study**

| **Primer** | **Sequence (5’-3’)** | **Source** |
|---|---|---|
| *Nde*I-*dmfase*-for | CTTTAAGAAGGAGATATA**CATATG**ACTGAAGCCAGCGAATCCTG | This study |
| *Hind*III-*dmfase*-rev | CTCGAGTGCGGCCGC**AAGCTT**GACCCGCGGCGCCGGCTCATC | This study |
| *Nde*I-*dmfase*-α2-for | CTTTAAGAAGGAGATATA**CATATG**AAAGACATTGCCATTCGCGGA | This study |
| *Hind*III-*dmfase*-α2-rev | TCGAGTGCGGCCGC**AAGCTT**GACCCGCGGCGCCGGCTCATCCTTG | This study |
| *dmfase*-S395A-for | TGCCAGCACGCTGGCCTATCTCGCCTATGCCAATG | This study |
| *dmfase*-S395A-rev | ATAGGCGAGATAGGCCAGCGTGCTGGCAATGAC | This study |
| *dmfase*-Y399A-for | GAGCTATCTCGCCGCTGCCAATGAGCAGATCATG | This study |
| *dmfase*-Y399A-rev | TCTGCTCATTGGCAGCGGCGAGATAGCTCAGCGTG | This study |
| *dmfase*-Y440A-for | ACGGGCTGTCAACCGCTGATGGTCATATCGACGGTC | This study |
| *dmfase*-Y440A-rev | CGATATGACCATCAGCGGTTGACAGCCCGTAATAG | This study |
| *dmfase*-H519A-for | TGCTGACCGGCTCGGCTCCCGAATATCAGACTTG | This study |
| *dmfase*-H519A-rev | TCTGATATTCGGGAGCCGAGCCGGTCAGCACAAC | This study |
| *dmfase*-E521A-for | CGGCTCGCATCCCGCATATCAGACTTGGGCCAATG | This study |
| *dmfase*-E521A-rev | CCAAGTCTGATATGCGGGATGCGAGCCGGTCAG | This study |
